# Cytokinin-inducible *DIRIGENT13* involved in lignan synthesis and ROS accumulation promotes root growth and abiotic stress tolerance in *Arabidopsis*

**DOI:** 10.1101/2024.06.25.600713

**Authors:** Alesia Melnikava, François Perreau, Candelas Paniagua, Sy-Chyi Cheng, Li-Min Huang, Elena Ubogoeva, Elena Zemlyanskaya, Yu-Liang Yang, Grégory Mouille, Dominique Arnaud, Jan Hejatko

## Abstract

Dirigent (DIR) proteins mediate regio- and stereoselectivity of phenoxy radical coupling of monolignols for (neo)lignan or lignin synthesis. However, the role of DIR proteins and lignans in plant development and response to environmental stresses remains elusive. Here, we provide a functional characterization of the cytokinin-responsive *DIRIGENT13* (*DIR13*) in *Arabidopsis thaliana*. DIR13 was localized to the root endodermis and at the margins of lateral root primordia and lateral roots. While not necessary for Casparian strip formation, DIR13 promoted both main and lateral root growth. Untargeted metabolomics and imaging mass spectrometry analyses unveiled the role of DIR13 in facilitating (neo)lignan synthesis in primary and lateral roots. Interestingly, DIR13 activated the production of putative oxomatairesinol and matairesinol-cysteine which are oxidative derivatives of the lignan matairesinol. Our data also identified DIR13 as an enhancer of salt and drought tolerance. Particularly, *DIR13* attenuated salt stress-mediated inhibition of germination and root growth. Moreover, DIR13 activated reactive oxygen species (ROS) accumulation both under control and salt stress conditions, and cytokinin further enhanced the salt-induced ROS production specifically in the *DIR13* overexpressing line. Our results uncover a role for DIR13-produced lignans in the priming of ROS accumulation, mediating both abiotic stress tolerance and the control of root architecture.

## Introduction

As sessile organisms, plants experience abiotic stress caused by multiple environmental factors, such as drought, salinity, and osmotic stress, which in turn induce alterations in plant physiological and biochemical processes, leading to compromised plant growth and developmental modifications (Zhao *et al*., 2020). Salt stress represents a critical environmental challenge with profound implications for crop growth and productivity, thereby leading to significant economic repercussions in agriculture (Qadir *et al*., 2014). Salt stress can detrimentally reduce photosynthesis, and impede seed germination and root growth (Zhao *et al*., 2020; Zou, Zhang and Testerink, 2022; Balasubramaniam *et al*., 2023).

Reactive oxygen species (ROS) play crucial roles in plant development and stress responses (Mittler, 2017; Mhamdi and Van Breusegem, 2018; Smirnoff and Arnaud, 2019). Various (a)biotic stresses induce an increase in ROS production in plants and hydrogen peroxide (H_2_O_2_) is particularly implicated in stress sensing, signaling, and adaptation (Miller *et al*., 2010; Smirnoff and Arnaud, 2019). However, excessive ROS generation under stresses leads to direct oxidative damage of DNA, lipids and proteins (Smirnoff and Arnaud, 2019). Besides their role in stress response, ROS also function as signaling molecules for optimal root growth and development (Tsukagoshi, 2016; Eljebbawi *et al*., 2020; Mase and Tsukagoshi, 2021). Basal ROS level and localized production are essential for cell expansion, root hair growth, lateral root development, cell polarity, and Casparian strip (CS) formation (Takeda *et al*., 2008; Lee *et al*., 2013; Manzano *et al*., 2014; Mangano *et al*., 2017). Although ROS have the ability to modulate root growth to some degree autonomously, they also engage in complex crosstalk with other signaling molecules including hormones (Mase and Tsukagoshi, 2021). However, despite its central importance, our knowledge of the mechanisms controlling localized ROS production in plants is still in its infancy.

The phenolic compounds lignans and neolignans are found throughout the plant kingdom and are thought to play roles in various physiological processes, including response to environmental stresses [(Paniagua *et al*., 2017) and references there in]. Lignan biosynthesis starts with the dimerization/radical coupling of two coniferyl alcohol molecules from the phenylpropanoid and monolignol pathways leading to (+/-)-pinoresinol formation (Davin *et al*., 1997; Suzuki and Umezawa, 2007). The term lignan refers to a class of phenolic secondary metabolites linked by an 8-8’ bond, while alternative dimers from 8-O-4’ and 8-5’ coupling are known as neolignans (Moss, 2000; Umezawa, 2003). Through a series of enzymatic reactions involving pinoresinol/lariciresinol reductase (PLR) and secoisolariciresinol dehydrogenase, pinoresinol can be converted *via* lariciresinol to secoisolariciresinol and matairesinol (Davin and Lewis, 2003; Suzuki and Umezawa, 2007). A variety of lignans can be derived from pinoresinol, lariciresinol, secoisolariciresinol, and matairesinol through aromatic substituent modification, such as glycosylation, hydroxylation, methylation, and cysteine addition (Umezawa, 2003; Niculaes *et al*., 2014). A number of pharmacological studies, mostly in mammal cells, showed that many (neo)lignans exhibit potent antioxidant properties (Zálešák, Bon and Pospíšil, 2019) while others have prooxidant activities (Hwang *et al*., 2012; Ko *et al*., 2021). Surprisingly, their roles in ROS metabolism in plants are poorly documented. It was suggested that the poplar phenylcoumaran benzylic ether reductase (PCBER) and related enzymes, such as PLRs, reduce (neo)lignans *in planta* to form antioxidants for protection against oxidative damage (Niculaes *et al*., 2014).

The first dirigent (DIR) protein was found to be responsible for the regio- and stereo-selective specificity of coniferyl alcohol dimerization during (+)-pinoresinol formation (Davin *et al*., 1997; Gang *et al*., 1999). DIRs do not possess oxidizing or any other enzymatic activity but rely on laccases (LAC) or apoplastic peroxidases (PER) to generate phenoxy radicals, which are then guided *via* oriented binding to DIR trimers towards regio- and enantioselective product formation (Davin *et al*., 1997; Halls *et al*., 2004; Gasper *et al*., 2016; Paniagua *et al*., 2017). DIR and DIR-like proteins have been identified in almost all vascular plants and were classified into six subfamilies DIR-a to DIR-f (Ralph *et al*., 2006; Ralph *et al*., 2007). In Arabidopsis, the DIR family consists of 25 members, but for most of them, their physiological functions remain unknown. Within the DIR-a subfamily, DIR5 and DIR6 participate in (-)-pinoresinol biosynthesis (Kim *et al*., 2012), and DIR12/DP1 is specific to the outer seed coat and essential for neolignans biosynthesis (Yonekura-Sakakibara *et al*., 2021). ESB1/DIR10, crucial for CS positioning, and DIR9, DIR16, DIR21, and DIR24 (DIR-e subfamily) were shown to be responsible for the majority of lignin deposition during CS formation (Lee *et al*., 2013; Gao *et al*., 2023). DIR13 (At4G11190) is the closest paralog of DIR5, DIR6, DIR14, and DIR12 but lacks the conserved residues necessary for (-)-pinoresinol formation and exhibits strong, although not exclusive, expression in the root (Kim *et al*., 2012). *DIR* genes are induced by biotic and abiotic stresses (Paniagua *et al*., 2017) and overexpression of some *DIRs* can lead to improved tolerance to biotic and abiotic stresses (dos Santos *et al*., 2022; Li *et al*., 2022).

Cytokinins are plant growth regulators controlling many important developmental and growth processes such as cell division and differentiation (Kieber and Schaller, 2018; Yang *et al*., 2021; Svolacchia and Sabatini, 2023). Cytokinins and the cytokinin signaling pathway are also involved in regulating biotic and abiotic stress response (Cortleven *et al*., 2019; Skalak *et al*., 2021). Several studies have demonstrated that cytokinins can exert both beneficial and detrimental effects on stress tolerance or resistance (Nishiyama *et al*., 2011; Reguera *et al*., 2013; Arnaud *et al*., 2017). The transcription factors type-B ARABIDOPSIS RESPONSE REGULATORs (ARRs) ARR1, ARR10, and ARR12 transcriptionally regulate a number of cytokinin-responsive genes (Taniguchi *et al*., 2007; Bhargava *et al*., 2013; Zubo *et al*., 2017; Xie *et al*., 2018), but the functional importance of the majority of type-B ARR targets remains to be clarified.

In this paper, we describe the spatiotemporal specificity of *DIR13* expression and DIR13 protein localization in the root and demonstrate that *DIR13* is under control of cytokinins and type-B ARRs. DIR13 activated the accumulation of several putative lignans and neolignans in the roots and primed ROS accumulation leading to improved plant adaptation to salinity and drought, and modifications of root architecture toward growth promotion. These observations imply a link between DIR13-mediated lignan production, ROS signaling, and regulation of plant development and stress acclimation.

## Results

### The root-expressed *DIR13* is upregulated by cytokinins

To determine precisely the localization of DIR13 in root cells, we generated Arabidopsis lines expressing the *pDIR13:NLS-3xGFP* construct. In the early (3-day-old) seedling development, we observed the activity of *DIR13* promoter predominantly in the root, covering the region ranging approximately from the root-shoot junction up to the root transition zone (Supplemental Figures 1A, B). Weak expression was detectable also in the shoot apical meristem (Supplemental Figure 1C). In the older (7-day-old) seedlings, *DIR13* promoter was active dominantly in the endodermis, but also in the epidermis, cortex, and pericycle of the root transition zone (Figure 1A), as well as in cells surrounding the lateral root (LR) primordia (Figure 1B).

**Figure 1.**
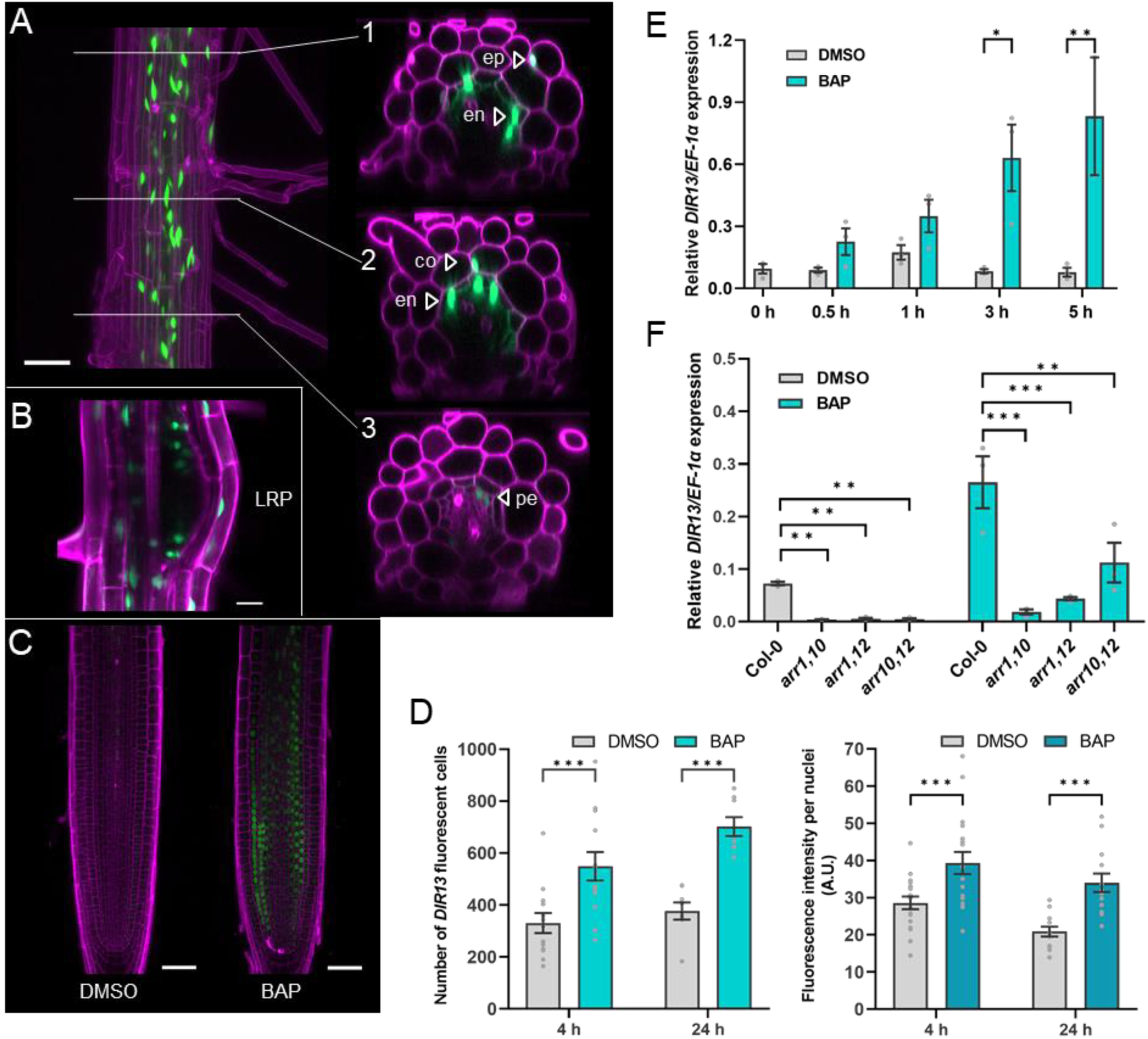
Cytokinin-inducible *DIR13* is active in all root tissues including lateral root primordia. **(A)** Representative images of the root differentiation zone from 7-day-old seedlings expressing *pDIR13:NLS-3xGFP* and radial optical cross-sections of the root cells (1–3). The fluorescent signal was detected in all layers of *Arabidopsis* root. **(B)** 9-day-old seedlings show the activity of *DIR13* promoter in lateral root primordia. In (A and B), the plasma membrane signal from PI staining is shown in magenta and GFP is in green. Scale bars represent 50 μm in (A) and 20 μm in (B). ep, epidermis; en, endodermis; co, cortex; pe, pericycle; LRP, lateral root primordia. **(C)** The promoter of *DIR13* is activated by cytokinin in the root apical meristem (RAM). Representative images of 5-day-old roots expressing *pDIR13:NLS-3xGFP* after 24h of 5 µM BAP and control (0.1% DMSO) treatments. The plasma membrane signal from PI staining is shown in magenta and GFP is in green. Scale bars represent 50 µm. **(D)** Left panel - number of fluorescent nuclei per root at the RAM of 5-day-old seedlings expressing *pDIR13:NLS-3xGFP* after 4h and 24h of 5 µM BAP and control (0.1% DMSO) treatments. Data are presented as the mean +/- SE (n ≥ 10). Asterisks indicate statistically significant differences between BAP and control treatments based on Poisson mixed model analysis and Tukey test (\*\**P* < 0.01, \*\*\**P* < 0.001). Right panel - the fluorescence intensity of individual nuclei after 4h and 24h of 5 µM BAP and control (0.1% DMSO) treatments was quantified in RAM. Results are the means +/- SE (n ≥ 10). Asterisks indicate statistically significant differences between BAP and control treatments based on two-way ANOVA followed by Tukey’s HSD test (\*\**P* < 0.01, \*\*\**P* < 0.001). **(E)** *DIR13* expression is inducible by cytokinin. *DIR13* expression was analyzed by RT-qPCR in roots of 11-day-old seedlings after 5 μM BAP and control (0.1% DMSO) treatments for 0, 0.5, 1, 3, and 5 hours. **(F)** *DIR13* expression is regulated by type-B ARR transcription factors. The expression level of *DIR13* was examined by RT-qPCR in *arr1,10, arr1,12* and *arr10,12* double mutants after 5 μM BAP and control (0.1% DMSO) treatments for 1h. In (C-D), transcript levels were normalized to the reference *EF-1α* gene, and the relative *DIR13/EF-1α* expression ratio is shown. Data are the means of a representative experiment. Error bars indicate +/- SE (n = 3). Asterisks indicate statistically significant differences between BAP and control treatments (C) or between genotypes (D) based on two-way ANOVA followed by Tukey’s HSD test (\**P* < 0.05, \*\**P* < 0.01, \*\*\**P* < 0.001).

*DIR13* expression was proposed to be controlled by cytokinins (Taniguchi *et al*., 2007; Bhargava *et al*., 2013). Accordingly, we observed that exogenous cytokinin application, 6-Benzylaminopurine (BAP) for 4 or 24 hours, significantly enhanced *DIR13* promoter activity in the root apical meristem (RAM) where only a weak signal was detectable under control conditions (Figures 1C, D). These observations were confirmed by RT-qPCR analyses showing a prompt upregulation of *DIR13* transcripts in roots from 3 hours after cytokinin treatment (Figure 1E). In the *DIR13* promoter, we found the presence of binding regions recognized by the type-B response regulators *ARR1, ARR10* and *ARR12* involved in cytokinin signaling [(Xie *et al*., 2018); Supplemental Figure 1D]. In line with that, we observed a significant decrease in *DIR13* expression in single *arr1-3, arr10-5* and *arr12-1* mutants compared to Col-0 WT (Supplemental Figure 1E). Even a more pronounced decrease in *DIR13* transcript levels was detectable in double *arr1,10, arr10,12* and *arr1,12* mutant backgrounds, both in the presence or absence of exogenous cytokinin (Figure 1F). In summary, our findings provide evidence for root-dominant expression of *DIR13* being under positive control of cytokinin signaling.

### DIR13 protein localizes to the root endodermis and periphery of developing lateral roots

Using *pDIR13:DIR13-mCherry* Arabidopsis lines, we found DIR13 being localized to the endodermis of the root differentiation zone at a domain resembling the Casparian strip (CS) (Figure 2A–C). However, compared to CASP1-GFP and ESB1-mCherry (Supplemental Figure 2), which are known to determine the CS positioning and regulate CS formation (Roppolo *et al*., 2011; Hosmani *et al*., 2013), DIR13 localization pattern was more diffuse and of weaker signal intensity. That might be due to the weak binding of DIR13 to the cell wall and possible wash-out during specimen preparation (see Materials and Methods). Following 24 h of BAP treatment, the DIR13 signal was observed earlier (more distally) in the root differentiation zone compared to control-treated samples (Figure 2D–E). Besides the endodermis of the main root, a strong DIR13 signal was also recognizable at the periphery of developing lateral roots, forming a ring-like distribution surrounding the lateral root primordia (LRP) and persisting there up to the lateral root (LR) emergence (Figure 2F–I). Therefore, the DIR13 localization to the root endodermis and peripheral cells of LRs suggests involvement of DIR13 in CS and/or LR formation.

**Figure 2.**
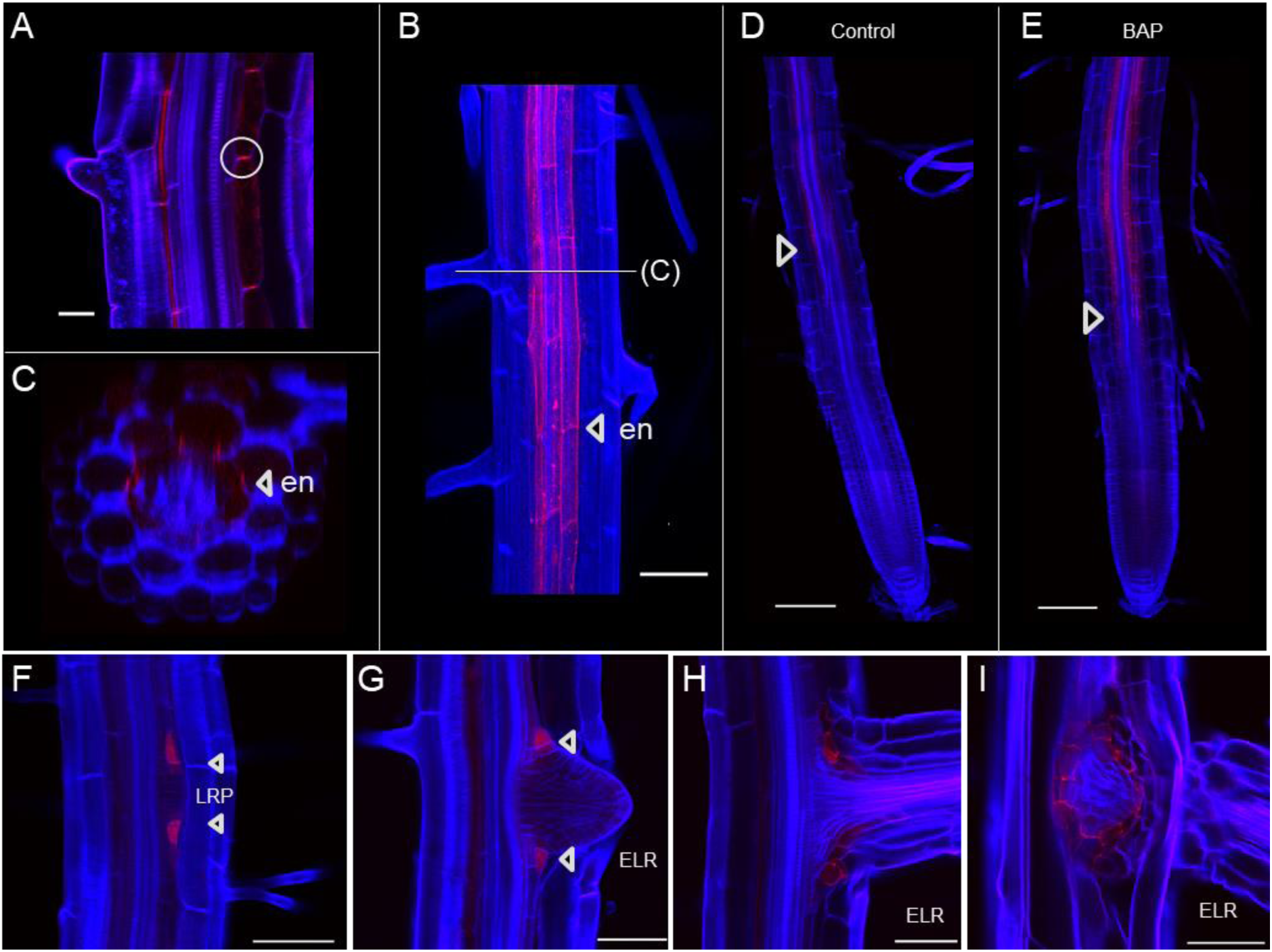
DIR13 localizes in the root endodermis and peripheral cells of lateral root primordia. **(A-C)** DIR13-mCherry localizes in the endodermal cells of the root differentiation zone. Representative confocal images of roots from 7-day-old seedlings expressing *pDIR13:DIR13-mCherry.* Median section (A), Z-stack overview (B), and radial optical cross-section (C) of the root. **(D, E)** Cytokinin treatment enhances DIR13-mCherry expression in roots. Representative Z-stack confocal images of 7-day-old seedlings after 24h of 5 µM BAP and control (0.1% DMSO) treatments. BAP shifts DIR13 localization closer to the root apical meristem, indicated by arrowheads. **(F-I)** DIR13-mCherry protein is detected in the peripheral cells of lateral root (LR) primordia (F) and emerged lateral root (G-I). Representative confocal images of different stages of LR development in 9-day-old seedlings. In (A-I), cleared roots were stained with Calcofluor White, the cell wall is shown in blue and mCherry is in red. Scale bars represent 50 μm in (A–C) and (F–I), and 100 μm in (D, E). en, endodermis; LRP, lateral root primordia; ELR, emerged lateral root.

### DIR13 does not interfere with Casparian strip formation

To explore the function of *DIR13* in root growth and development, we produced the *DIR13* overexpressing (OE) lines *35S:DIR13* #6 and #8 with expression levels about 33 and 85 times higher than WT respectively (Supplemental Figure 3A). Because only knock-down T-DNA insertion mutants (*dir13-1*, *dir13-2*, and *dir13-3*) were available, we used CRISPR-Cas9 genome editing to generate a knock-out *dir13-4* mutant producing truncated *DIR13* transcripts at position 128 of the coding sequence (Supplemental Figure 3B-D). Importantly, the expression level of the closest *DIR13* homologs *DIR6* and *DIR14,* as well as *DIR10 (ESB1),* was not affected in *DIR13* OE and *dir13-4* mutant lines (Supplemental Figure 3 E), *DIR5* and *DIR12* being not expressed in the root.

According to previous reports (Roppolo *et al*., 2011; Hosmani *et al*., 2013), the examination of lignin autofluorescence indicated a complete loss of well-organized CS structure leading to a patchy and disrupted morphology in mutants deficient in *ESB1* or *CASP1/3* (Supplemental Figure 4). In contrast, no visible changes in the structure of CS were detectable in *DIR13* deficient *dir13-4* or *DIR13* OE lines when compared to Col-0 WT (Supplemental Figure 4A–F). We performed propidium iodide (PI) staining to assay any potential delay in the formation of a CS-mediated apoplastic barrier (Supplemental Figure 4G–N). We confirmed previous findings (Roppolo *et al*., 2011; Hosmani *et al*., 2013) demonstrating a delayed CS development in *casp1-1;3-1* and *esb1-1* mutants with an extended penetrability of PI into the central cylinder of about 15 endodermal cells in comparison to Col-0 WT (Supplemental Figure 4L–O). In the *dir13-4* mutant and *DIR13* OE line, no significant delay in CS formation was observed (Supplemental Figure 4G–K, O). Overall, our data suggest that in contrast to ESB1/DIR10, DIR13 is not essential for CS formation and function.

### DIR13 controls root architecture

In the developing lateral roots, DIR13-mCherry, CASP1-GFP and ESB1-mCherry exhibited distinct localization patterns. While CASP1 was nearly absent in the LRP (Supplemental Figure 5A–C), ESB1 localization is restricted to the endodermal cells of newly formed LR, contrasting with the localization of DIR13 to the peripheral cells surrounding the LRP and emerged LR (Supplemental Figure 5D–K).

To determine whether DIR13 contributes to root growth and development, we examined root phenotypes of 9-, 11- and 15-day-old *dir13-4* and *DIR13* OE seedlings. In *DIR13* OE lines, the lateral root growth was enhanced as indicated by a higher number of emerged LRs, and an increase in LR density and total LR length (Figure 3A, C–D). In addition, the LR density and total LR length were decreased in the *dir13-4* mutant, although the reduced number of emerged LRs remained not statistically significant when compared to Col-0 WT (Figure 3B, F–H). Besides the positive effects on the LR growth, the overexpression of *DIR13* also increased the length of primary roots and the RAM size (higher number of meristematic cells) compared to WT (Supplemental Figures 5L-Q). However, no changes in primary root growth or RAM size were detected for the *dir13-4* mutant. Altogether, our findings indicate that DIR13 controls root architecture by promoting lateral root formation and growth. And when overexpressed, DIR13 has the potential to enhance the main root growth.

**Figure 3.**
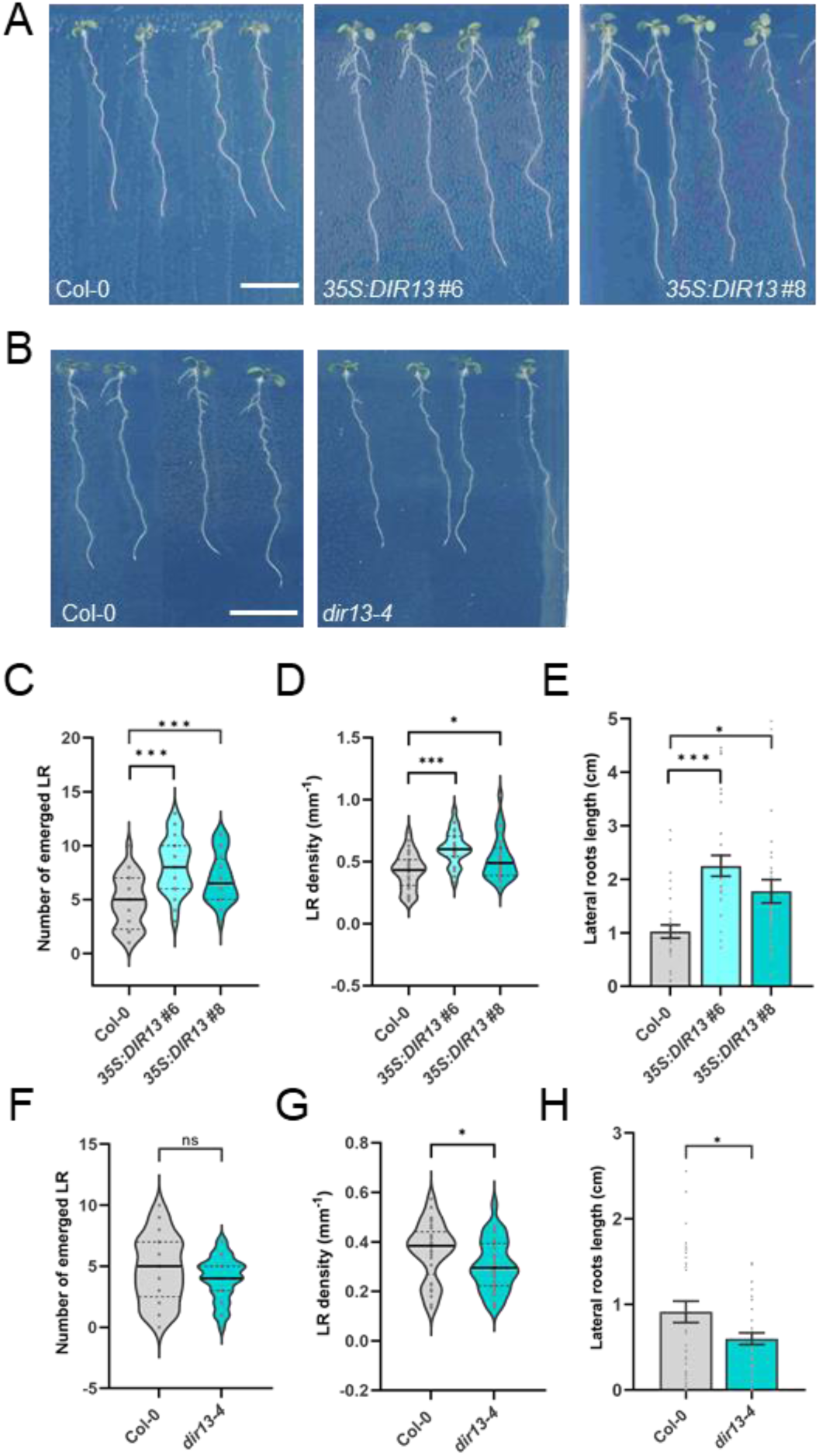
DIR13 promotes lateral root growth. **(A)** The phenotype of 11-day-old Col-0 WT, *DIR13* overexpressing lines (*35S:DIR13* #6 and #8) and **(B)** *dir13-4* knock-out mutant seedlings growing under normal conditions. Scale bars represent 1 cm. **(C, D, F**, and **G**) Average number of emerged lateral roots (C and F) and LR density expressed as the number of lateral roots per mm of primary root length (D and G) in 11-day-old Col-0 WT, *35S:DIR13* #6 and #8 (C and D) and *dir13-4* mutant (F and G) seedlings. In (B, C, F, and G), data show the median and upper and lower quartiles of a representative experiment (n ≥ 30). Asterisks indicate statistically significant differences between genotypes at \**P* < 0.05, \*\**P* < 0.01 and \*\*\**P* < 0.001 based on Poisson mixed model analysis (C and F), one-way ANOVA followed by Tukey’s HSD test (D), and unpaired t-test (G). (**E** and **H**) Total lateral root length per seedling (in cm) in 11-day-old Col-0 WT, *DIR13* overexpressing (E) and *dir13-4* mutant (H) seedlings. In (E and H), data are the means of a representative experiment. Error bars indicate +/- SE (n ≥ 30). Asterisks indicate statistically significant differences between genotypes at \**P* < 0.05, \*\**P* < 0.01 and \*\*\**P* < 0.001 based on one-way ANOVA followed by Tukey’s HSD test (D) and unpaired t-test (H).

### DIR13 improves salt stress tolerance

Considering the potential role of DIRIGENTs in plant response to environmental constraints (Paniagua *et al*., 2017; dos Santos *et al*., 2022; Li *et al*., 2022), we inspected the possible role of DIR13 in the salt stress response. When placed on control media (0 mM NaCl), the germination rate of the *DIR13* OE or *DIR13*-deficient lines was similar to Col-0 WT (Supplemental Figure 6A, D). In the presence of 150 mM NaCl, the germination rates of *DIR13* OE and *dir13-4* seeds were higher and lower, respectively, than in Col-0 WT (Figure 4A, B), and at 50 and 100 mM NaCl, the *DIR13* OE lines also showed improved germination rate (Supplemental Figure 6B–C, E–F).

**Figure 4.**
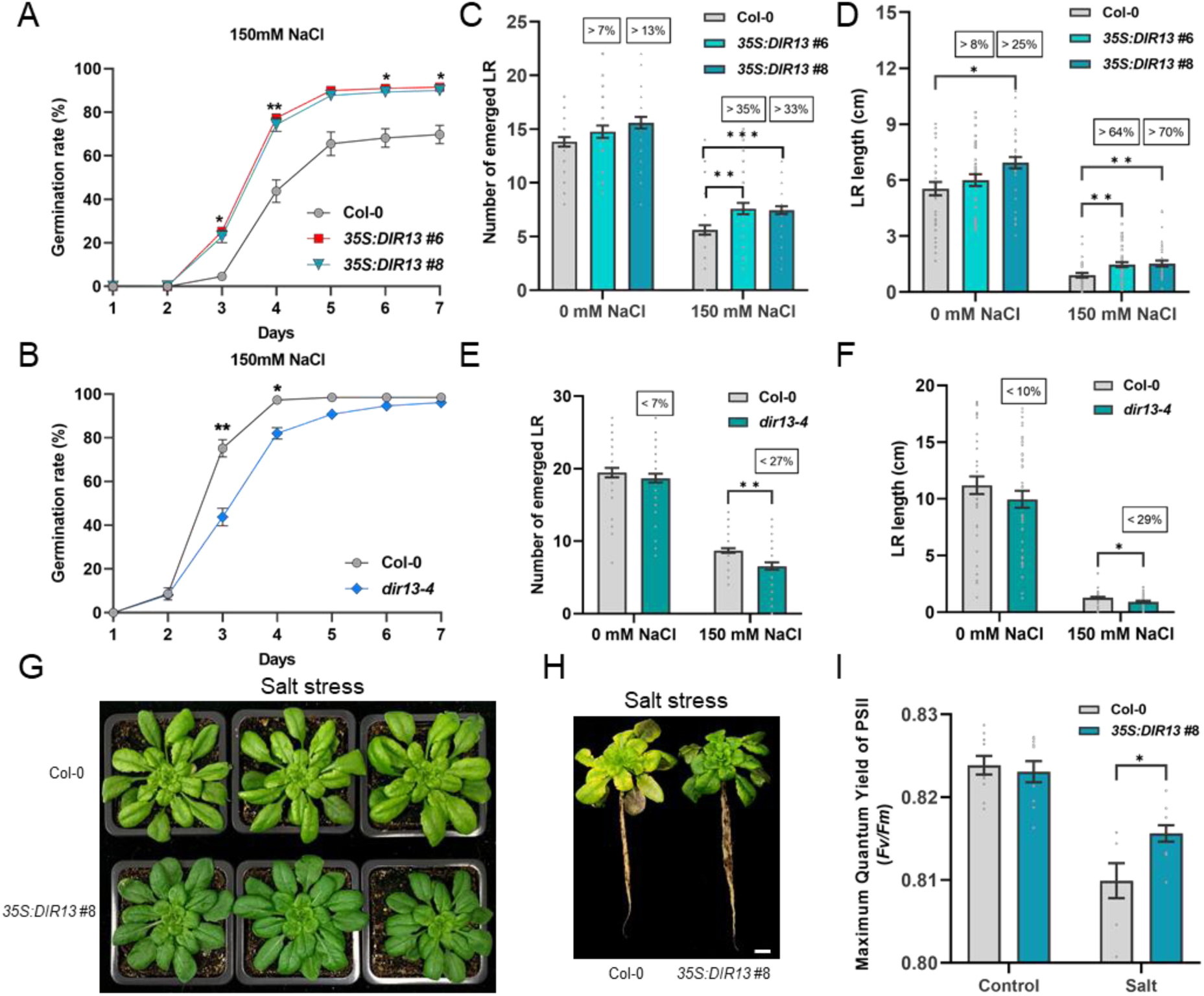
*DIR13* improves plant tolerance to salt stress. **(A, B)** Seed germination rates of Col-0 WT, *35S:DIR13* #6 and #8 overexpressing lines (A) and *dir13-4* mutant (B) under salt stress treatment. The germination rate was recorded for 7 days after stratification in the presence of 150 mM NaCl (for 0, 50 and 100 mM NaCl treatments see Suppl. Fig. 6). Data are the averages of three individual experiments. Error bars indicate +/- SE, (n ≥ 100). Asterisks indicate statistically significant differences between Col-0 WT and *DIR13* OE in (A) and between Col-0 WT and *dir13-4* in (B) based on repeated measures two-way ANOVA followed by Dunnett’s HSD test (\**P* < 0.05, \*\**P* < 0.01). **(C-F)** DIR13 promotes lateral root growth in the presence of salt. The average number of emerged lateral roots (C, E) and total lateral root length in cm per seedling (D, F) were measured 9 days after the transfer of Col-0 WT, *35S:DIR13* #6, #8 (C-D) and *dir13-4* (E-F) seedlings to 150 mM NaCl. Data show the means of a representative experiment +/- SE (n ≥ 38). Asterisks indicate statistically significant differences between genotypes at \**P* < 0.05, \*\**P* < 0.01 and \*\*\**P* < 0.001 based on Poisson mixed model analysis (C and E), and two-way ANOVA followed by Tukey’s HSD test (D and F). Numbers in rectangles show the relative difference compared to Col-0 in percentage. **(G, H)** Phenotypes of 8-week-old Col-0 WT and *35S:DIR13* #8 plants growing in short-day conditions after 4 weeks of progressive salt stress by application of increasing NaCl concentrations (100, 150, 200, and 300 mM) each week. Scale bar represents 1 cm. **(I)** Quantification of Photosystem II maximum quantum yield efficiency (*Fv*/*Fm*) of Col-0 WT and *35S:DIR13* #8 plants after 23 days of progressive salt stress. Data are the averages of a representative experiment. Error bars indicate +/- SE, (n ≥ 7). Asterisks indicate statistically significant differences between Col-0 WT and *35S:DIR13* #8 based on two-way ANOVA followed by Tukey’s HSD test (\**P* < 0.05).

The effects of salt stress on root growth were examined after transferring 5-day-old seedlings to 150 mM NaCl-supplemented media. The *DIR13* OE lines were more resistant to NaCl-induced inhibition of root growth with a significant increase in primary root elongation, number of emerged LR, LR density, and LR length compared to Col-0 WT (Figure 4C–D; Supplemental Figure 6G, I). Conversely, the *dir13-4* mutant was more sensitive than WT to salt stress for all these root phenotypes (Figure 4E–F; Supplemental Figure 6H–J). Importantly, the observed relative differences in the increased or decreased root growth between the *DIR13* OE or *dir13-4* lines respectively, and Col-0 WT were more pronounced upon salt treatment than in control condition (Figure 4C-F). Therefore, besides promoting root growth, DIR13 also improves tolerance to salt stress.

We have also inspected the role of DIR13 in the response to progressive salt stress by watering 4-week-old plants grown on soil with increasing NaCl concentrations every week (see Materials and Methods). In the absence of stress (water control), the growth and development of Col-0 WT and *DIR13* OE were indistinguishable (Supplemental Figure 7A-B). After 4 weeks of salt stress, a similar reduction in rosette area was observed for Col-0 WT and *DIR13* OE lines (Supplemental Figure 7B). However, while Col-0 WT plants exhibited a striking yellowish leaf phenotype and defects in the root growth, the *DIR13* OE plants remained green, revealing higher dark-adapted quantum efficiency of photosystem II (*Fv/Fm*) and enhanced root system when compared to WT controls, suggesting improved stress tolerance (Figure 4G– I). Altogether, these findings suggest that *DIR13* promotes salt stress tolerance throughout plant development from seed germination to adult plants.

### Plant drought tolerance is enhanced when *DIR13* is overexpressed

To further investigate DIR13 function in plant response to abiotic stresses, we performed drought stress experiments on 4-week-old Col-0 WT and *DIR13* OE plants exposed to 3 weeks of water withholding. The plants were then rewatered for recovery. Compared to Col-0 WT, *DIR13* OE plants exhibited higher tolerance to drought stress, as evidenced by a lower percentage of wilted plants (Supplemental Figure 7C) and a greater survival rate after 5 days of recovery (Supplemental Figure 7D, E). *DIR13* OE plants also showed larger shoot area during the drought stress stage compared to Col-0 WT plants which was even more notable during the recovery phase (Supplemental Figure 7F). Additionally, higher *Fv/Fm* was detected in drought-stressed *DIR13* OE compared to Col-0 WT plants, indicating a better PSII efficiency (Supplemental Figure 7G). Collectively, these data suggest that DIR13 positively regulates acclimation to abiotic stresses and may play an important role in plant responses to environmental challenges.

### DIR13 activates ROS accumulation in roots

Salt stress triggers the accumulation of ROS which act as an important signaling messenger or a toxic by-product of stress responses (Miller *et al*., 2010). To further clarify the role of DIR13 in response to salt treatment (150 mM NaCl for 30 min), in the presence and absence of cytokinin (BAP), we analyzed the level of intracellular ROS in the cortex and epidermis of the root differentiation zone using the ROS fluorescent dye 2’,7’-dichlorodihydrofluorescein-diacetate (H_2_DCFDA; Figure 5). Interestingly, in control conditions, the basal ROS level in roots was significantly higher (1.3 times) in *DIR13* OE line and lower (0.67 times) in *dir13-4* mutant than in Col-0 WT. Moreover, NaCl-triggered ROS accumulation was more pronounced in *DIR13* OE and reduced in *dir13-4* compared to WT. Pretreatment with BAP did not change the ROS levels in all the lines when compared to the control condition. However, when combined with salt stress, BAP further enhanced the salt-induced ROS production only in the *DIR13* OE line (Figure 5). In summary, DIR13 promotes ROS accumulation under normal and salt stress conditions, and cytokinin enhances salt-mediated ROS production when *DIR13* is overexpressed.

**Figure 5.**
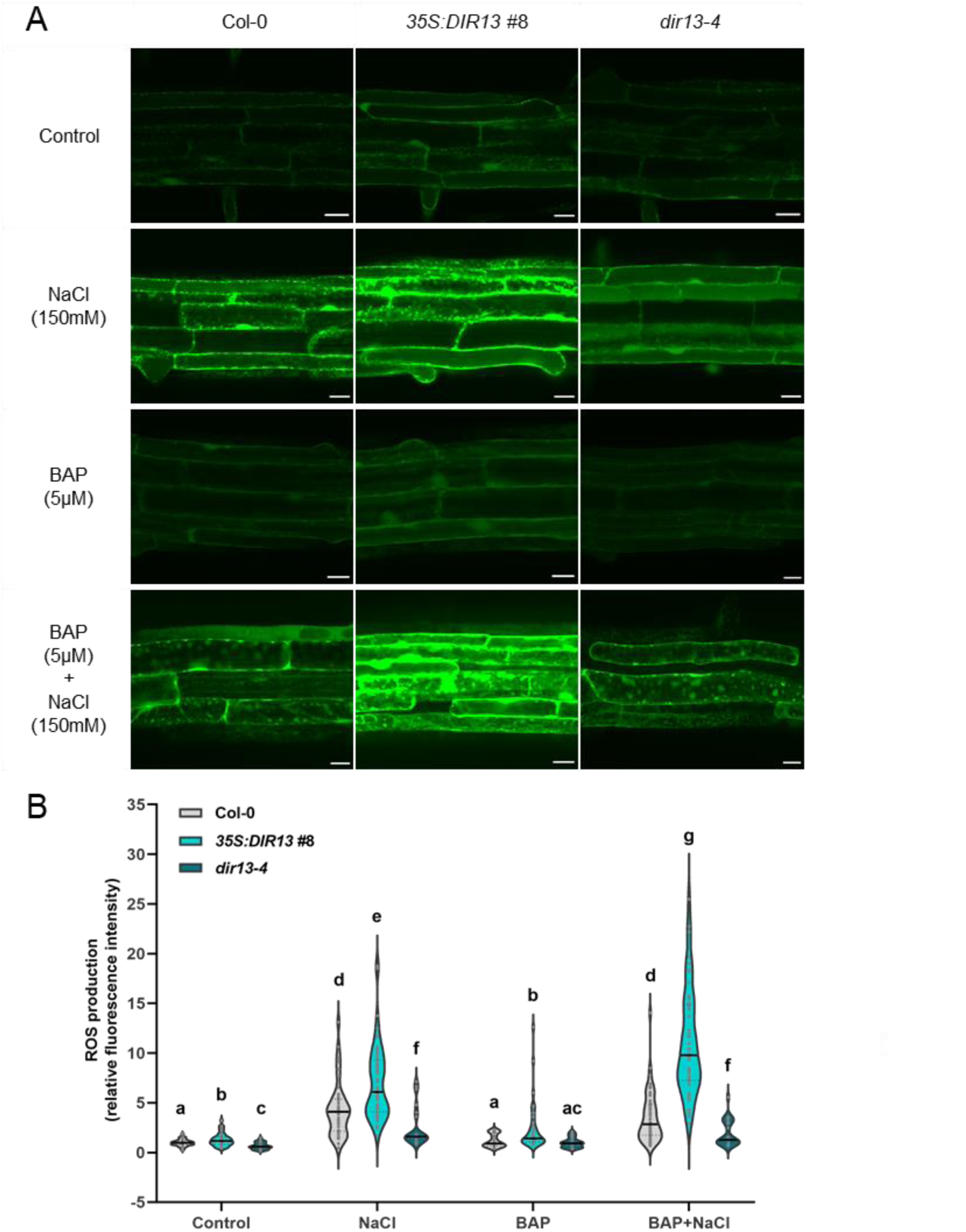
Salt stress-induced ROS accumulation in root cells is promoted by *DIR13*. **(A)** Representative confocal images and **(B)** quantification of ROS production measured with the fluorescent dye H_2_DCFDA in the root differentiation zone of 7-day-old Col-0 WT, *35S:DIR13* #8 and *dir13-4* seedlings pretreated for 2 hours with 5 µM BAP or 0.1% DMSO as control and exposed to 150 mM NaCl or control solution (0 mM NaCl) for 30 min. In (A), H_2_DCFDA fluorescence is shown in green and scale bars represent 20 μm. In (B), data show the median and upper and lower quartiles of three independent experiments (n ≥ 20). Different letters indicate statistically significant differences at *P* < 0.05 between genotypes and treatments based on repeated measures two-way ANOVA followed by Tukey’s HSD test.

The production of apoplastic ROS was also tested by luminol assays in leaves and roots after treatment with flg22, an epitope of the bacterial flagellin eliciting immune responses in plants (Felix *et al*., 1999). Albeit statistically not significant, flg22-triggered ROS burst was stronger in the leaves and particularly the roots of *DIR13* OE line when compared to Col-0 WT (Supplemental Figure 8A, B). In contrast, disease symptoms and bacteria content were similar between Col-0 WT and *DIR13* OE line 3 days after spray-inoculation with the virulent bacteria *Pseudomonas syringae* pv. tomato DC3000 (Supplemental Figure 8C, D). Based on these observations, it seems that DIR13 plays a preferential role in the response to abiotic stress. However, the positive role of DIR13 in pathogen-elicited ROS production cannot be excluded.

### DIR13 promotes the production of putative (neo)lignans promoting lateral root formation

It was shown that DIR12, a close homolog of DIR13, is involved in neolignan biosynthesis (Yonekura-Sakakibara *et al*., 2021). To provide more insights into the possible role of DIR13 in (neo)lignan production, two untargeted metabolome analyses of root extracts from 10-day-old *DIR13* OE lines or *dir13-4* mutant were performed by HPLC-MS/MS together with Col-0 WT as a control. For the experiment with *DIR13* OE lines, 1221 and 1382 compounds were detected in ElectroSpray Ionization in positive (ESI+) and negative (ESI-) modes, respectively (Supplemental Table 1). For the *dir13-4* mutant, 1214 and 872 compounds were identified in ESI+ and ESI-modes (Supplemental Table 2). Pairwise comparisons with Col-0 WT identified 143 (ESI+) and 225 (ESI-) differentially accumulated metabolic features (DAMF) at *P* ≤ 0.05 in the *DIR13* OE lines, and 36 (ESI+) and 26 (ESI-) DAMF in the *dir13-4* mutant. Principal component analysis and the heatmaps built using correlation distance and average linkage illustrated the differences in metabolite accumulation within tested genotypes (Figure 6A; Supplemental Figures 9A, B and 10A).

**Figure 6.**
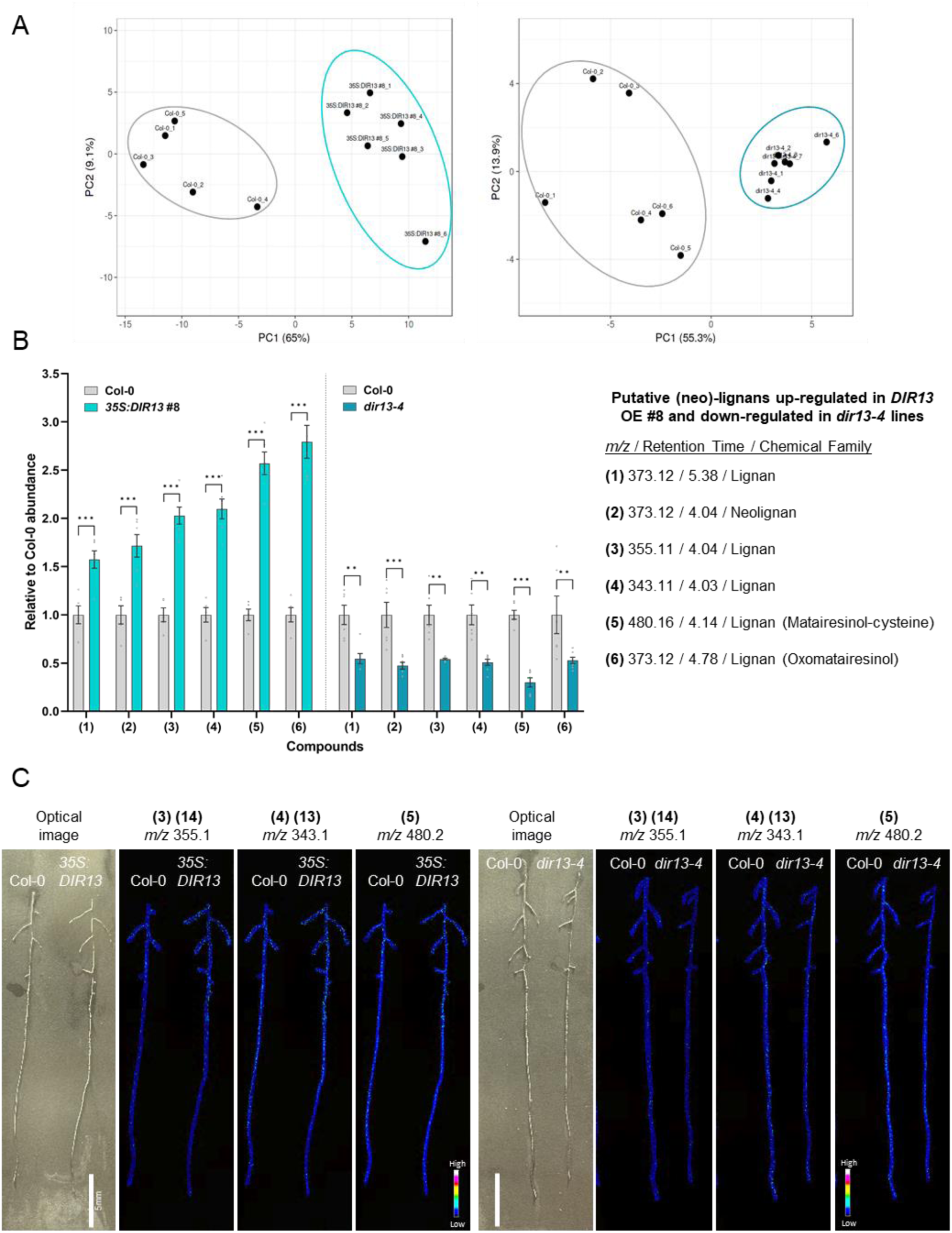
DIR13 mediates (neo)lignan formation in planta. **(A)** Principal Component Analysis (PCA) of HPLC-MS/MS data in ElectroSpray Ionization in positive (ESI+) mode from root biological replicates of 10-day-old Col-0 WT and *35S:DIR13* #8 (left panel) and Col-0 WT and *dir13-4* (right panel). PCA demonstrates the difference genotypes in the accumulation of significantly different abundant compounds. **(B)** *DIR13* overexpression leads to the accumulation of putative lignans and neolignans in roots. Selected differentially abundant compounds annotated as putative (neo)-lignans were extracted from Col-0 WT, *35S:DIR13* #8 and *dir13-4* roots and detected by HPLC-MS/MS in ESI+ mode. Data are shown as relative to Col-0 WT and are the means +/- SE (n ≥ 5). Asterisks indicate statistically significant differences between Col-0 WT and *35S:DIR13* #8, and Col-0 WT and *dir13-4* based on two-way ANOVA followed by Tukey’s HSD test. (\**P* < 0.05, \*\**P* < 0.01, \*\*\**P* < 0.001). **(C)** Spatial distribution of putative lignans in the roots of *DIR13* overexpressing and *dir13-4* mutant lines. Imaging Mass Spectrometry (IMS) of roots from 10-day-old Col-0 WT, *35S:DIR13* #8 overexpressing and *dir13-4* mutant lines. Signals at *m/z* 355.1 (3) (14), *m/z* 343.1 (4) (13), *m/z* 480.2 (5) correspond to putative matairesinol (5), and metabolites annotated as lignans (3), (4), (13) and (14). The spatial resolution is 50 µm. MS signal intensity is indicated by color bars and the scale bar represents 5 mm. The experiment was repeated at least twice with similar results.

The majority of identified compounds had no annotation in homemade or public libraries. Metabolites with ≥1.5- or ≤0.5-fold change in accumulation between *DIR13* OE or *dir13-4* and Col-0 WT and with the highest significant differences at *P* ≤ 0.01 were selected for annotations (see Materials and Methods). Interestingly, for *DIR13* OE line in ESI+ mode, among 16 annotated compounds, 12 of them were putative lignans and neolignans, such as matairesinol-cysteine (5a; compound # is shown in parenthesis in the further text, see also Supplemental Figure 9C and Table 1), oxomatairesinol (6) and lariciresinol 4-O-glucoside (11), two of them were annotated as flavonoids [2-arylbenzofuran (8) and kaempferol-hexoside-rhamnoside (9)], and compound (7) corresponded to a fatty acid. Matairesinol (5b) was considered as an in-source fragment of matairesinol-cysteine (5a) suggesting that matairesinol is mainly oxidized by cysteine addition in Arabidopsis roots. Most of the putative (neo)lignans [compounds (1) to (7) and (12) to (16)] were up-regulated in *DIR13* OE compared to Col-0 WT except the monolignol glucoside syringin (10) and the lignan lariciresinol 4-O-glucoside (11) that were down-regulated (Supplemental Figure 9C). For the *dir13-4* mutant, among nine DAMF selected for identification in ESI+ mode, six compounds (1)-(6) were annotated as putative lignans or neolignans and were also identified in *DIR13* OE (Supplemental Figure 10B and Table 2). Compared to Col-0 WT, these six putative (neo)lignans were over-accumulated in *DIR13* OE but down-regulated in *dir13-4* mutant (Figure 6B and Supplemental Figure 13). The putative fatty acid compound (7) and flavonoid compounds (8) and (9), were also commonly identified in both overexpressing and mutant lines, where compounds (7) and (8) had an opposite abundance between *DIR13* OE and *dir13-4* and compound (9) was down-regulated in both *DIR13* OE and *dir13-4* roots (Supplemental Figures 9C and 10B). Surprisingly, in ESI-mode, putative (neo)lignans were not detected among 31 DAMF selected for annotation in *DIR13* OE lines (Supplemental Table 1), while in *dir13-4* mutant, out of 16 metabolites selected for identification in ESI-, four corresponded to (neo)lignans (Supplemental Table 2). In addition, no significant difference in the accumulation of anthocyanins between Col-0 WT and *DIR13* OE lines was observed (Supplemental Figure 9D).

To test the possible role of lignans in the control of root architecture, Arabidopsis seedlings were grown in the presence of matairesinol. Interestingly, we observed that when applied at low concentrations (1 and 10 nM), matairesinol promoted both LR density and the main root elongation (Supplemental Figure 12). This result suggests that matairesinol and possibly other (neo)lignans upregulated by DIR13 may have the potential to promote root growth.

Collectively, our data suggest that DIR13 promotes (neo)lignans synthesis with consequences on the accumulation of other phenolic compounds in Arabidopsis roots. Furthermore, the DIR13-mediated upregulation of matairesinol and possibly other (neo)lignans seems to have the potential to control root growth.

### DIR13-mediated accumulation of (neo)lignans in diverse root areas

Imaging mass spectrometry (IMS) has been widely used to analyze the spatial abundance of metabolites in plant tissues (Boughton *et al*., 2016). We developed a new matrix-assisted laser desorption/ionization (MALDI)-IMS method with a spatial resolution of 50 µm by using carboxymethylcellulose (CMC) to stabilize 11-day-old Arabidopsis roots on the sample target (see Materials and Methods). By employing this approach, we examined the spatial distribution of (neo)lignans along the root in *DIR13* OE and *dir13-4* lines. For the imaging, we selected six putative (neo)lignans in ESI+ mode that were both up-regulated in *DIR13* OE line and down-regulated in *dir13-4* mutant [compounds (1) to (6) (Figure 6 B)] or only up-regulated in *DIR13* OE line [compounds (12 to 16) (Supplemental Figure 9 C)]. Many ions identified as putative (neo)lignans by LC-MS/MS have similar mass-to-charge (*m/z*) ratios that cannot be distinguished by IMS with a mass range of ± 0.1 Da (Figure 6C and Supplemental Figure 11). Nevertheless, in meristematic and mature areas of the main root and in the lateral roots (Figure 6C), we were able to detect signals in positive-ion mode at *m/z* 355.1 related to lignans (3) and (14), *m/z* 343.1 for putative lignans (4) and (13), and *m/z* 480.2 corresponding to matairesinol-cysteine (5a). Compared to Col-0 WT, these putative lignans (at *m/z* 355.1, 343.1 and 480.2) were more abundant in *DIR13* OE line, preferentially in the mature portion of the main root while in the *dir13-4* mutant, the signal of these ions was weaker in all root areas.

No apparent difference in the distribution of the ion at *m/z* 373.1 corresponding to putative (neo)lignans (1), (2), (6) and syringin (10) was detected between Col-0 WT, *DIR13* OE line and *dir13-4* mutant (Supplemental Figure 11), possibly because syringin was found to be down-regulated in *DIR13* overexpressing lines by LC-MS/MS analysis (Supplemental Figure 9C). The ions at *m/z* 359.1 and 583.2 probably related to putative matairesinol (5b) and the neolignan (15) respectively, were only downregulated in the *dir13-4* mutant root compared to Col-0 WT (Supplemental Figure 11). Even if ionization by NH4^+^ is unlikely to occur during IMS, the signals of putative lignan (12) at *m/z* 512.2 and neolignan (16) at *m/z* 556.2 from possible ionization by NH4^+^ or at *m/z* 494.1 (12) and 539.2 (16) from ionization by H^+^ indicate that these ions might also be up- and down-regulated in the roots of *DIR13* OE and *dir13-4* lines, respectively (Supplemental Figure 11). Altogether, the IMS results support our LC-MS/MS metabolomic data and strongly suggest that DIR13 promotes the accumulation of (neo)lignans in diverse areas of Arabidopsis root.

## Discussion

### Does DIR13-mediated lignan production control ROS formation?

Our data from the MS-based metabolomic profiling and imaging approaches demonstrate that DIR13 is involved in the (neo)lignan production that correlates with increased ROS production. The positive impact of DIR13 on the ROS accumulation can be seen under control conditions, but particularly in the salt stress-induced roots. Stress-induced apoplastic ROS production is dominantly mediated via peroxidases (PERs) and NADPH oxidases (NOXs), the latter referred to as respiratory burst oxidase homologs (RBOHs) in plants [(Suzuki *et al*., 2011; Chapman *et al*., 2019; Smirnoff and Arnaud, 2019; Castro *et al*., 2021) and references there in].

The ability of (neo)lignans to impact ROS homeostasis was demonstrated previously. On one hand, lignans and neolignans, including DIR13-upregulated matairesinol, are known antioxidants and ROS scavengers (Lee *et al*., 2012; Zálešák *et al*., 2019). On the other hand, several lignans are able to upregulate ROS production. For instance, (+)-medioresinol mediates its antifungal activities via inducing intracellular ROS accumulation in *Candida albicans* (Hwang *et al*., 2012). Gomisin L1, a lignan isolated from *Schisandra berries*, induces apoptotic cell death by upregulating intracellular ROS production via activating NOX activity (Ko *et al*., 2021).

Another mechanism of DIR13-mediated ROS regulation can be associated with lignan biosynthesis. Synthesis of (neo)lignans leads to the consumption of H_2_O_2_ or O_2_ if the monolignol substrates are oxidized by apoplastic peroxidases (PERs) or laccases (LAC), respectively (Davin *et al*., 1997). It was suggested that DIR12/DP1, a close homolog of DIR13, works with LAC5 for neo(lignan) synthesis in seeds (Yonekura-Sakakibara *et al*., 2021). If this is the case also for DIR13, then oxygen will be consumed for oxidizing monolignols leaving H_2_O_2_ (not used by PERs) accumulating in the apoplast. Moreover, PER enzymes have a dual function depending on the biological context. PERs consume H_2_O_2_ through the peroxidase cycle for cell wall cross-linking, lignan synthesis or lignification, or produce ROS through the oxidative cycle upon stress perception (Bolwell *et al*., 2002; Smirnoff and Arnaud, 2019). PERs substrates are phenolic compounds and our metabolomic analysis indicates a decrease in putative syringin, a glycosylated monolignol, in *DIR13* OE lines. It is therefore possible that DIR13-mediated (neo)lignan accumulation induces changes in PER activity and subsequently ROS accumulation. This scenario is further supported by the occurrence of an oxomatairesinol and matairesinol-cysteine, putative oxidized forms of the lignan matairesinol, that we found to be up- and down-regulated in *DIR13 OE* and *dir13-4* mutant respectively. Formation of cysteine adducts of coniferyl alcohol by PER activity was observed *in vitro* in the presence of H_2_O_2_ (Cong *et al*., 2013) and cysteine dilignols accumulated in the poplar with attenuated ROS scavenging capacity (Niculaes *et al*., 2014). However, whether some of the DIR13-synthesized lignans contribute to the regulation of the activity of ROS-producing enzymes, remains to be identified.

### DIR13 is involved in the control of root growth

ROS were demonstrated as potent regulators of root architecture, controlling both main root growth and lateral root formation (Mase & Tsukagoshi, 2021). For the regulatory role of ROS, its precise localization and chemical nature seems to play an important role. RBOHs catalyze the reduction of oxygen to superoxide (O_2_^•−−^), which rapidly dismutates to H_2_O_2_. Besides their role in lignan and lignin formation, a reaction consuming H_2_O_2_, PER can also produce ROS (O_2_^•−^, OH^•^ and H_2_O_2_) depending on the chemical environment (Smirnoff and Arnaud, 2019). In the main root, a differential distribution of H_2_O_2_ and superoxide along the root axis was observed (Dunand, Crèvecoeur and Penel, 2007). Superoxide may stimulate cell division in the RAM, while H_2_O_2_ might promote cell differentiation (Tsukagoshi, Busch and Benfey, 2010). Several PERs are implicated in the control of main root growth via regulating ROS distribution in the RAM (Passardi *et al*., 2006; Tsukagoshi, Busch and Benfey, 2010). The highly localized ROS production is also necessary for the CS formation in the endodermis, where RBHOF with the possible contribution of RBHOD were shown to be the main ROS-producing enzymes (Lee, *et al*., 2013; Fujita *et al*., 2020).

ROS also regulates LR formation. A number of *PERs* were found to be expressed during LR formation, and both H_2_O_2_ and superoxide accumulate in the LRP (Manzano *et al*., 2014). Notably, *per7* and *per57* mutants have a lower number of LRs and reduced main root length. MYB DOMAIN PROTEIN 36 (MYB36)-regulated PRX9/64 might also be important for LR emergence (Fernández-Marcos *et al*., 2017). H_2_O_2_ was found to accumulate in the middle lamellae of cortical and endodermal cells overlying newly formed LRPs and several RBOHs were found to be expressed in the developing LRP, particularly at their edges (Orman-Ligeza *et al*., 2016). It was suggested the role of RBOHs in facilitating the LR emergence via ROS-mediated cell wall remodeling in the overlying tissues (Orman-Ligeza *et al*., 2016). However, other studies indicate that RBOHD/F inhibit LR development (Ma *et al*., 2012; Li *et al*., 2015). The *rbohD*/*F* mutant exhibits increased LR density that was correlated with an increase in PER activity and superoxide accumulation in the mature root zone (Li *et al*., 2015).

Our results showed that *DIR13*, expressed in the endodermis of the main root and at the margin of LRs, is necessary for main root growth and LR emergence and growth. While we cannot exclude a direct role of (neo)lignans in root growth independently of ROS, it is likely that, DIR13-regulated ROS accumulation (due to its ROS scavenging and/or altered activity of ROS-producing enzymes) regulates both main root and LR growth. The highly specific localization of DIR13 supports this type of regulation. In the main root, DIR13 overlaps with the CS domain in the endodermis and localizes to the outskirts of LRP, i.e., in the tissues where the functional importance of ROS homeostasis in the control over the main root and LR formation, respectively, was demonstrated. This is well in line with tight spatial control over ROS production/homeostasis and its functional importance in plant development and stress response (Castro *et al*., 2021; Mase and Tsukagoshi, 2021). The absence of DIR13 does not prevent the plant from forming CS or LRs. In the case of CS, this may be attributed to the functional redundancy with other DIRs, as recently demonstrated for DIR9, DIR16, DIR21, and DIR24, shown to be responsible for the majority of lignin formation in the CS (Gao *et al*., 2023). Nonetheless, rather than being directly involved in the cell wall lignification, DIR13-mediated lignan production seems to provide an additional regulatory mechanism, fine-tuning the ROS-regulated control over the root architecture, facilitating thus the plant’s accommodation to physiological and/or developmental status.

### DIR13-induced ROS production facilitates salt and drought adaptation

Our results clearly show that DIR13 plays an important role in salt and drought stress tolerance for different assays including seed germination, root growth and progressive stress to soil-grown plants. This is in agreement with previous findings based on expression analyses, suggesting the role of dirigent genes in various stress responses (Paniagua *et al*., 2017; Luo *et al*., 2022). For instance, recent studies have shown that silencing of pepper *CaDIR7* weakened plant defense and increased susceptibility to biotic and salt stresses (Khan *et al*., 2018). Another example is the overexpression of sugarcane *ScDIR* genes in *Nicotiana benthamiana* leading to enhanced drought tolerance (Li *et al*., 2022). However, the precise mechanism of DIRs action in stress adaptation remains unknown.

Induced ROS production as a signaling molecule and activation of antioxidant machinery to scavenge harmful ROS have an essential role in the response and adaptation to salt, one of the most severe abiotic stresses in plants (Miller *et al*., 2010; Kesawat *et al*., 2023). We showed that DIR13 activates a rapid (1 h) overaccumulation of ROS upon salt treatment. An initial increase in apoplastic ROS mediated by NADPH oxidases or PERs is one of the mechanisms by which plants respond to (a)biotic stress (Smirnoff and Arnaud, 2019). PERs participate in plant resistance to pathogens but their precise role in response to abiotic stresses remains unclear. Nevertheless, both RBOHD and RBOHF play a crucial role in salt stress tolerance (Jiang *et al*., 2012; Ma *et al*., 2012) and RBOHD/F-dependent ROS production is required for salt-induced antioxidant defense and ROS-scavenging enzymatic activities in Arabidopsis (Ben Rejeb *et al*., 2015). Besides being the key mechanism for the local stress response, ROS production also mediates the systemic acquired acclimation, the response induced by abiotic stress and activating resistance or acclimation pathways in tissues that were not yet subjected to the stress [reviewed in (Castro *et al*., 2021; Kesawat *et al*., 2023; Mittler & Blumwald, 2015; Wang *et al*., 2024)].

The observation that *DIR13* overexpression improved plant tolerance to long-term salt treatment suggests a better protection against oxidative damage. Considering our data implying the positive role of DIR13 in the ROS production under both control and stress conditions, we hypothesize that DIR13-mediated upregulation of ROS levels is a signal to induce higher salt and drought tolerance, probably via an H_2_O_2_-priming mechanism (Hossain *et al*., 2015).

### Cytokinin-controlled DIR13 provides a possible link between hormonal- and ROS-mediated regulations

We showed that both basal expression of *DIR13* and its upregulation by exogenous cytokinins are dependent on the presence of functional RRBs ARR1, ARR10 and ARR12 transcription factors, activating cytokinin signaling (Sakai *et al*., 2001; Argyros *et al*., 2008). Like DIR13, cytokinins were found to regulate root growth and stress adaptation. Cytokinins are potent regulators of the root architecture with contrasting effects depending on root developmental areas (Ramireddy, Chang and Schmülling, 2014; Yamoune *et al*., 2021). On one hand, cytokinins promote root growth by stimulating stem cell activity (Higuchi *et al*., 2004; Nishimura *et al*., 2004; Köllmer *et al*., 2014). On the other one, cytokinins restrict root growth by inducing the differentiation of the dividing cells which results in a reduction of the RAM size (Dello Ioio *et al*., 2007, 2008; Street *et al*., 2016; Di Mambro *et al*., 2019). Cytokinins also inhibit LR initiation and early development (Miyawaki *et al*., 2006; Kuderová *et al*., 2008; Laplaze *et al*., 2008; Chang, Ramireddy and Schmülling, 2013) but seem to promote LR elongation at later stages (Rani Debi, Taketa and Ichii, 2005; Li *et al*., 2006). The ROS-mediated regulation of root growth was shown to be both auxin- and cytokinin-independent (Tsukagoshi, Busch and Benfey, 2010). However, the cytokinin kinetin induces the accumulation of two lignans (phyllanthin and hypophyllanthin) in *Phyllanthus amarus* shoot cultures (Sparzak-Stefanowska and Krauze-Baranowska, 2022). Therefore, cytokinin-regulated (neo)lignan production through DIR13 might connect the ROS- and hormonal-mediated regulations of root architecture.

Cytokinin signaling generally inhibits numerous stress responses including salt and drought stresses with some exceptions showing positive effects on plant tolerance to environmental constraints (Cortleven *et al*., 2019; Skalak *et al*., 2021). One mechanism could be the cytokinin-mediated ROS production (Novák *et al*., 2013; Wang *et al*., 2015; Arnaud *et al*., 2017; Nongpiur *et al*., 2024). Cytokinin-induced ROS production through ARR2-mediated transcriptional regulation of several apoplastic peroxidases (e.g. *PER33/34*) promotes Arabidopsis resistance to *Pseudomonas syringae* bacteria (Arnaud *et al*., 2017). Moreover, an increase in endogenous cytokinin levels by induction of *ISOPENTENYL TRANSFERTASE* (*IPT*) expression in tobacco (*Nicotiana tabacum*) induces ROS accumulation in leaves (Novák *et al*., 2013). The overexpression of Arabidopsis *IPT8* leads to higher ROS production only under salt stress condition (Wang *et al*., 2015). This was correlated with transcriptional upregulation of *RBOH* genes and downregulation of genes for ROS scavenging enzymes including *PERs*. Altogether, this suggests that changes in the cytokinin homeostasis may sensitize or prime the plant to the salt stressor, leading to higher ROS production (Hossain *et al*., 2015; Cortleven *et al*., 2019). A similar mechanism, possibly including synergistic effects of DIR13-mediated lignan production and exogenously applied cytokinin can explain the cytokinin-upregulated ROS production specifically in the *DIR13* OE lines. Nonetheless, the detailed molecular mechanisms underlying the cytokinin-dependent lignan-mediated regulation of ROS production and its developmental importance remain to be identified.

### Model for the DIR13-regulated ROS production and root growth control

Our findings demonstrate the novel role of the DIR13 protein in the regulation of ROS accumulation, root architecture and stress adaptation. We propose that in response to different developmental and/or environmental signals, *DIR13* is activated, at least in part via the cytokinin signaling pathway, inducing (neo)lignan production in roots. This leads to ROS accumulation, helping the plant to fine-tune the root architecture under stress conditions which are nearly permanently present in nature (Figure 7). Noteworthy, DIR13-mediated root growth and the potential benefits of (neo)lignans for plant stress tolerance as well as human health (Adlercreutz, 2007) make DIR13 and possibly other DIRs an attractive breeding target.

**Figure 7.**
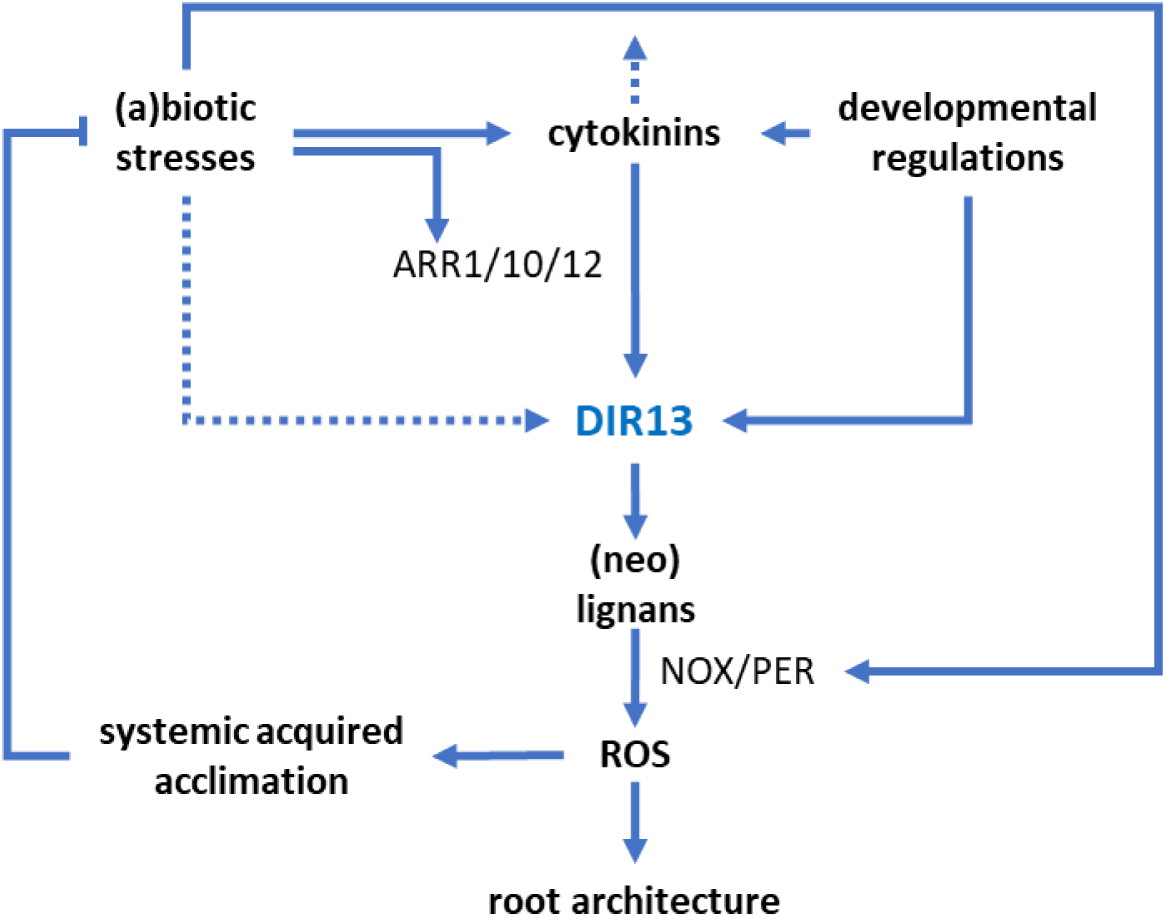
Proposed mechanism of DIR13 action in the abiotic stress response. In a response to different developmental and/or environmental signals, *DIR13* is activated, at least partially via the cytokinin-regulated multistep phosphorelay (MSP) signaling. (A)biotic stresses may activate the MSP either via controlling the abundance of endogenous cytokinins and/or by affecting some of the numerous stress-related MSP inputs e.g. ABA signaling (Skalak et al., 2021) affecting directly the type-B response regulators. DIR13-mediated (neo)lignans accumulation upregulates NOX/PER-mediated ROS production. ROS act as effector molecules modulating the stress adaptive response by inducing systemic acquired acclimation and root architecture. The ROS production induced by abiotic stress seems to be sensitized by cytokinins via unknown mechanism.

## Materials and Methods

### Plant material and growth conditions

The mutant and reporter lines used in this study are in *Arabidopsis thaliana* Col-0 background. *dir13-1* (N838075), *dir13-2* (N808825 in Col-3 background), *dir13-3* (N565236) were obtained from the Nottingham Arabidopsis Stock Centre and T-DNA insertions were confirmed by PCR genotyping with primers listed in Supplemental Table 3. T-DNA insertion sites for the *dir13* mutants were determined by sequencing. The type-B ARR mutants *arr1-3* (N6971), *arr2-5* (N425801), *arr10-5* (N39989), *arr11-3* (N506544), *arr12-1* (N6978), *arr1-3 arr10-5* (N39990), *arr1-3 arr12-1* (N6981), and *arr10-5 arr12-1* (N39991) were previously described (Mason *et al*., 2005; Argyros *et al*., 2008; Hill *et al*., 2013). *CASP1-GFP* and *ESB1-mCherry* lines used for Casparian strip visualization were generated and described by Roppolo *et al*. (2011) and Hosmani *et al*. (2013) respectively. Mutants *casp1-1;3-1* and *esb1-1* were previously described by Baxter *et al*. (2009) and Roppolo *et al*. (2011).

Surface sterilized seeds were sown on half-strength Murashige–Skoog (1/2 MS) medium (Duchefa) MES-KOH pH 5.7 (Duchefa), and 1 % (w/v) plant agar (Duchefa)and stratified for 2 days at 4°C in the dark. Seedlings were grown under long-day conditions (16 hours light at 21°C/8 hours dark at 19°C) with the light intensity at 130 µmol m^−2^ s^−1^.

For cytokinin treatment followed by either confocal microscopy or gene expression analysis, 6 or 11-day-old seedlings were transferred in 2 ml of liquid 1/2 MS media (pH 5.7) and incubated for 2 h in the growth chamber for recovery. Then, 6-Benzylaminopurine (BAP, Duchefa) at a final concentration of 5 µM or 0.1 % DMSO as a control were added and seedlings were further incubated in the growth chamber for 1 to 24 hours.

For drought and salt stress experiments on adult plants, stratified seeds were cultivated on soil in a growth chamber under short-day conditions (10 hours light at 22 °C / 14 hours dark at 19 °C) with a relative humidity of 40 - 60 % and the light intensity at 130 µmol m^−2^ s^−1^.

### Cloning and generation of transgenic plants

To generate *DIR13* promoter fusion line with GFP (*pDIR13:NLS-3xGFP*) and *DIR13* overexpression lines (*35S:DIR13)*, we applied Ligase Independent Cloning (LIC) according to (de Rybel *et al*., 2011). To produce the *pDIR13:NLS-3xGFP* reporter construct, 3.1 Kb of *DIR13* promoter was PCR-amplified using Phusion DNA Polymerase (Thermo Fisher Scientific) from Col-0 gDNA. The amplified fragment, including LIC adapter sites, was introduced into the pPLV04 LIC-based vector containing 3 repeats of Green Fluorescent Protein (3xGFP) fused to the nuclear localization signal (NLS). For the generation of *35S:DIR13* overexpression lines, the full-length coding sequence (CDS) of *DIR13* was PCR-amplified from Col-0 gDNA, including LIC adapter sites and the fragment was inserted into pPLV26 LIC-based vector with the 35S promoter from cauliflower mosaic virus. The primers with appropriate adapter sites used for the cloning are listed in Supplemental Table 3. To generate the translational fusion of DIR13 with mCherry (*pDIR13:DIR13-mCherry*), the region ∼3.1 Kb upstream of ATG along with the CDS of *DIR13* prior to the stop codon, was PCR-amplified using Herculase II Fusion DNA polymerase (Agilent Technologies) and primers containing PacI and SnaBI restriction sites (Supplemental Table 3). The PCR products were digested and cloned into the intermediate vector pZEO-mCherryT35S (Samalova *et al*., 2023), and then sub-cloned through the GATEWAY^TM^ LR reaction in the pFAST-G01 destination vector (Shimada, Shimada and Hara-Nishimura, 2010). The fidelity of the constructs was confirmed by sequencing. Arabidopsis Col-0 plants were transformed using the GV3101 strain of Agrobacterium tumefaciens according to the floral dip protocol (Clough and Bent, 1998). Transgenic lines were isolated on 1/2 MS media supplemented with appropriate antibiotics. Transgenic lines were screened for 3:1 segregation of the resistance marker and raised to homozygous T3 lines. 3, 3 and 5 independent lines were selected for further experiments for *pDIR13:NLS-3xGFP, 35S:DIR13* and *pDIR13:DIR13-mCherry,* respectively.

For the generation of the *dir13-4* knock-out mutant line, we applied CRISPR/Cas9 genome editing technology. The cloning was performed by the ProTech Facility/CRISPR lab at Vienna Biocenter Core Facilities GmbH (Richter *et al*., 2018). The vector was created using a Golden Gate-based cloning system to simultaneously target two locations with two single-guide RNAs (sgRNAs) in the *DIR13* gene and delete the coding sequence at position 128 from the start codon along with the 3’-UTR, leading to a null mutation of *DIR13* (Supplemental Table 3; Supplemental Figure 3). The final vector was transformed into *A. tumefaciens* GV3101 cells containing the pGreen helper plasmid pSOUP (Hellens *et al*., 2000). Agrobacterium-mediated transformation of Arabidopsis was performed as described above. T1 transgenic plants were selected with YFP fluorescent seed coat marker and grown in soil. The deletion event was confirmed by PCR amplification of the target sites and sequencing using primers specified in Supplemental Table 3. Potential heterozygous plants were propagated to T2 generation. T2 seeds lacking YFP fluorescence were grown in soil and screened by PCR to confirm the absence of Cas9 nuclease gene and the deletion of *DIR13* gene. 10 lines of T3 plants were screened to ensure homozygosity for the *dir13* mutation and 2 selected for further experiments. *DIR13* expression was checked by semi-quantitative PCR and RT-qPCR (Supplemental Table 3).

### Gene expression analysis

Total RNA was extracted from 11-day-old roots using Trizol reagent (Invitrogen). Genomic DNA (gDNA) contamination was removed by DNase I digestion (DNA-free; Thermo Fisher Scientific) according to the manufacturer’s protocol. First-strand cDNA was synthesized from 1 µg of total RNA using RevertAid First Strand cDNA Synthesis Kit and oligo (dT)_18_ primers (Thermo Fisher Scientific) and diluted 2 times prior to use. Real-time PCR was performed on the cDNAs using FastStart SYBR Green Master (Roche) with primers listed in Supplemental Table 3. Real-time quantification was made using the Rotor-Gene Q 72-slots system and analyzed with the Rotor-Gene Q Series Software (QIAGEN). The cycling conditions were as follows: an initiation denaturation step at 95 °C for 5 minutes, then 45 cycles of 30 seconds at 95 °C, 30 seconds at 60 °C, and 60 seconds at 72 °C. The amplified samples were normalized against the reference genes *Elongation factor 1α* (*EF-1α* At5G60390) and *Ubiquitin 1* (*UBQ1* At3G52590). For semi-quantitative PCR, the first-strand cDNA was diluted 10-fold and used as a template for PCR amplification (using Dream-Taq DNA polymerase (ThermoFisher), 10mM dNTPs and *EF-1α/DIR13* primers). The primers are specified in Supplemental Table 3.

### Confocal laser scanning microscopy and image analysis

To visualize the cell wall, propidium iodide (PI, Sigma) staining was done as described previously (Alassimone, Naseer and Geldner, 2010). Alternatively, seedlings cleared overnight in ClearSee solution were then stained in 0.1 % Calcofluor White (Sigma) according to (Ursache *et al*., 2018). For more details see Suppl. Materials and Methods. A Zeiss LSM 880 laser-scanning microscope with a 25X objective (LD LCI Plan-Apochromat, 0.8 Imm Korr DIC M27) was used to observe the fluorescence from GFP, mCherry, PI and Calcofluor White staining with the pinhole set to 1 airy unit. The excitation and emission wavelengths were set as follows: GFP, 488 and 490–550 nm; propidium iodide (PI), 561 and 595–700 nm. mCherry fluorescence was detected at a wavelength range 580-650 nm with 561-nm laser excitation. Calcofluor White staining was excited at 405-nm laser and detected at 425–475 nm to visualize the cell wall. Images were processed with ZEN2 (blue edition) and/or using Image J 1.53m. GFP fluorescence quantification of the *pDIR13:NLS-3xGFP* line was done using IMARIS 9.2 software (Bitplane, http://www.bitplane.com/imaris/imaris). The GFP and mCherry signal was confirmed in at least three independent lines.

### ROS localization and quantification in the root cells

To visualize the accumulation of intracellular ROS in the root differentiation zone, 2′,7′- dichlorofluorescein diacetate (H_2_DCFDA, Sigma) was used, following the protocol described by Achard *et al*. (2008) with some modifications. 6- to 7-day-old seedlings were transferred to 1/2 MS liquid media and placed back for 2 hours in the growth chamber for recovery. For cytokinin treatment, the seedlings were pre-treated with 5 µM BAP or 0.1 % DMSO (control) in liquid 1/2 MS media for 2 hours. The liquid media was then gently removed from the wells and replaced with a solution of 50 µM H_2_DCFDA (dissolved in liquid 1/2 MS media) and 150 mM NaCl or water as a control. The samples were incubated in the dark at 4 °C for 30 minutes. Seedlings were then washed with a solution containing 10 mM MES, 0.1 mM KCl, and 0.1 mM CaCl_2_ (pH 6.0) and incubated in liquid 1/2 MS media at room temperature for 30 minutes before microscopy. H_2_DCFDA fluorescence was observed with a Zeiss LSM 880 laser-scanning microscope after excitation at 488 nm and emission was detected at 500–550 nm with the pinhole set to 1 airy unit. Quantification of H_2_DCFDA fluorescence of the same regions of interest for each sample was performed using Image J 1.53m and results are shown in arbitrary units.

### Seed germination assay under salt stress treatment

Surface-sterilized seeds of *DIR13* overexpression and knock-out mutant lines with their respective wild-type controls from matched seed lots were sown on 1/2 MS agar plates supplemented with different concentrations of NaCl (0, 50, 100, or 150 mM) (Lachner). After stratification, the seed germination was monitored every 24 hours for 7 days and scored as the percentage of seeds with an obvious protrusion of the radicle through the seed coat.

### Root phenotyping

To analyze root architecture, Arabidopsis seedlings were grown vertically on MS/2 plates (under normal or salt-stressed conditions) and scanned after 9, 11 and 15 days of growth using a desktop scanner (Epson Perfection V700). Measurements of primary root length (in cm), average number of emerged lateral roots, lateral root density (in mm^-1^), and total lateral root length per seedling (in cm) were performed using ImageJ 1.53m. Primary root length was determined by measuring from the root tip to the base of the hypocotyl. The number of emerged lateral roots per seedling was determined by counting the visible lateral roots that penetrated the root epidermis. Lateral root density in the root branching zone was settled as the number of emerged lateral roots divided by the length of the primary root from the shoot base to the most rootward emerged LR. And, the total lateral root length was established by summing the length of all emerged lateral roots per seedling.

### Root growth assay under NaCl treatment *in vitro*

5-day-old plants vertically grown on agar plates were transferred to fresh plates supplemented with 0 and 150 mM NaCl. At the time of transfer, the position of the root cap was marked for each root. After 7 days, the length of the primary root was measured from the marked area to assess the primary root elongation under salt stress. For the measurements, the plates were scanned using a desktop scanner (Epson Perfection V700), and primary root growth, LR length, number, and density were quantified with Image J 1.53m software as described above.

### Salt stress assays

To evaluate the response of plants to progressive salt stress, 4-week-old plants initially grown on soil under normal conditions were exposed to an additional 4-week treatment with increasing concentrations of NaCl (100, 150, 200, and 300 mM) each week or water as control. The watering regime was adjusted to incorporate the NaCl solutions. Representative pictures were taken at the onset of salt stress symptoms (e.g. yellowing of the leaves). To assess the efficiency of Photosystem II maximum quantum yield (QY) under salt stress, chlorophyll fluorescence measurements were taken on the 23d day of salt stress using the automatic phenotyping platform (PlantScreen^TM^ SC Root System, Photon System Instrument (PSI). Maximum quantum yield (*Fv/Fm*) representing the maximal photochemical efficiency.

### Extraction of phenolic compounds from root tissue

Phenolic compounds were extracted from 10-day-old Arabidopsis roots using a modified protocol from (Routaboul *et al*., 2006). The samples were collected and ground in liquid nitrogen using the Qiagen Tissue Lyser for 2 minutes at 18 Hz with metal beads. To extract the phenolic compounds, the samples were homogenized in 1 mL cold acetonitrile/water (75:25, v/v) by vortexing for 1 minute and then sonicated on ice for 20 minutes. An internal standard, 1 µg of apigenin (Sigma), was added to the samples. After centrifugation at 16000 *g*, for 10 minutes at 4 °C, the pellet was further extracted with 1 ml acetonitrile/ water (75:25; v/v) overnight at 4 °C. The two extracts were pooled together, centrifuged again and the supernatants were used for HPLC-MS/MS analysis.

### HPLC–MS/MS analysis of phenolic compounds and data processing

The samples containing the phenolic compounds were dried down in a SpeedVac vacuum concentrator (o/n) and resuspended in 200 µl of ULC/MS grade water (Biosolve). Untargeted metabolomic data were acquired using a UHPLC system (Ultimate 3000 Thermo) coupled to a quadrupole time of flight mass spectrometer (Q-Tof Impact II Bruker Daltonics, Bremen, Germany).

A Nucleoshell RP 18 plus reversed-phase column (2 x 100 mm, 2.7 µm; Macherey-Nagel) was used for chromatographic separation with mobile phases (A) 0.1 % formic acid in H_2_O; and (B) 0.1 % formic acid in acetonitrile. The flow rate was 400 µL min^-1^. Data-dependent acquisition methods were used for mass spectrometer data in positive and negative ESI modes using the following parameters: capillary voltage, 4.5 kV; nebulizer gas flow, 2.1 bar; dry gas flow, 6 L min^-1^ ; drying gas in the heated electrospray source temperature, 140 °C. Samples were analyzed at 8 Hz with a mass range of 100–1500 *m/z*. Stepping acquisition parameters were created to improve the fragmentation profile with a collision RF from 200 to 700 Vpp, a transfer time from 20 to 70 µsec, and collision energy from 20 to 40 eV. Each cycle included a MS full scan and 5 MS/MS Collision Induced Dissociation on the 5 main ions of the previous MS spectrum. HPLC–MS/MS data processing was performed as described in Boutet *et al*. (2022).

### Metabolite annotation of untargeted metabolomic data

Metabolite annotation was performed as described previously by (Boutet *et al*., 2022). For differentially accumulated metabolites that had no or unclear annotation, we used SIRIUS 4.0 software (https://bio.informatik.uni-jena.de/software/sirius/) which proposes molecular structures using LC–MS/MS data (Djoumbou Feunang *et al*., 2016; Dührkop *et al*., 2021). Not annotated differentially accumulated metabolites that belong to molecular network clusters containing annotated metabolites were assigned to the same chemical family and their MS/MS spectra were tentatively annotated manually. Raw data were normalized with the internal standard (Apigenin) and the weight of the material used for the extraction. Metaboanalyst (https://www.metaboanalyst.ca/) and R software (R Core Team, https://www.R-project.org) were used for statistical analysis.

### Principal component analysis and heatmaps

Principal component analysis (PCA) and heatmaps were created by using ClustVis software (columns were centered, unit variance scaling was applied to columns, (https://biit.cs.ut.ee/clustvis/) (Metsalu and Vilo, 2015) based on the correlation distance and average linkage illustrating the difference between genotypes in the accumulation of significantly different abundant compounds.

### Imaging mass spectrometry analysis

To stabilize the roots on the stainless steel sample target, 100 µl carboxymethylcellulose (CMC, Sigma) 2 % (w/v) solution dissolved in water was spread and let dry evenly on a pre- heated target at 40 °C using a razor blade. After removal of the shoots, roots were delicately placed on the CMC-coated target and a flow of N_2_ gas was gently applied for root adhesion to the CMC-coated target. Universal MALDI matrix containing 2,5-dihydroxybenzoic acid (5 mg/mL) and α-cyano-4-hydroxycinnamic acid (5 mg/mL) dissolved in 50 % acetonitrile and 0.1 % trifluoroacetic acid was sprayed over the whole root-target with a HTX TM-Sprayer (one pass, 0.1 mL/min at 80°C, 10 psi N_2_ gas pressure; HTX Technologies LLC, USA). A mixture of peptide calibration standard (Bruker Daltonics) and universal matrix was added on the target edge and let dry for a few minutes for subsequent mass calibration. MALDI-IMS experiments were carried out with an autoflex maX MALDI-TOF/TOF mass spectrometer equipped with the Smartbeam-II laser (Bruker Daltonics, Bremen, Germany) operated by flexControl (version 3.4, Bruker Daltonics). Positive ion mass spectra were acquired in reflectron mode, in *m/z* 160– 1500 range with deflection of the ions below *m/z* 160. Laser settings were as follows: 333 Hz repetition rate, 75% intensity, ‘medium’ laser size corresponding to ∼45 μm diameter laser footprint. Mass spectrometer settings were as follows: ion source 1: 19.00 kV; ion source 2: 16.65 kV; lens: 8.35 kV; reflector: 21 kV; reflector 2: 9.40 kV; pulsed ion extraction: 150 nsec. The detector was set to 2190 V, and the digitizer was set to 5 GS sec^-1^. FlexImaging (version 3.0) was employed to visualize the spatial distribution of molecules in the sample. Each pixel in the MS image is related to a signal in a single mass spectrum acquired as a sum of 500 laser shots. The IMS experiments of roots were performed at a spatial resolution of 50 µm. The masses of CMC and CMC plus 1/2 MS media only were tested as negative controls.

### Statistical analysis

The experiments reported here were repeated at least three times with similar results unless otherwise mentioned. Statistical analyses were performed using GraphPad Prism version 9.4.1.681 for Windows, GraphPad Software, San Diego, California USA (www.graphpad.com). Unpaired t-test or one-way ANOVA followed by Tukey’s honestly significant difference (HSD) post hoc test were used. For pairwise comparisons and repeated experiments, 2-way ANOVA or mixed model ANOVA with random effects for different experiments were applied, followed by Tukey’s or Dunnett’s multiple comparisons tests. In the case of non-normal count data (e.g., number of cells), a Poisson mixed model was applied to identify differences between genotypes using R software (R Core Team, https://www.R-project.org). Significant differences for all statistical analyses were considered at *P* values ≤ 0.05 between compared samples.

## Author contributions

J.H. conceived the research and secured funding, A.M., F.P., C.M.P., S.-C. C., L.-M. H., E.Z., and D.A. performed the research, A.M, F.P., S.-C. C., Y.-L. Y., G.M., D.A. and J.H. analyzed and interpreted the data and A.M., D.A. and J.H. wrote the manuscript with the contribution of other co-authors.

## Acknowledgments

We are grateful to Niko Geldner for providing the CASP1-GFP, ESB1-mCherry, *esb1-1,* and *casp1-1;3-1* lines. We also thank Plant and Imaging (CELLIM) core facilities for helping with the growth of the plant material under controlled conditions and confocal microscopy. This work has benefited from the support of IJPB’s Plant Observatory platform P0-Chem.

## Supplemental Information

### Supplemental Figures

**Supplemental Figure 1.**
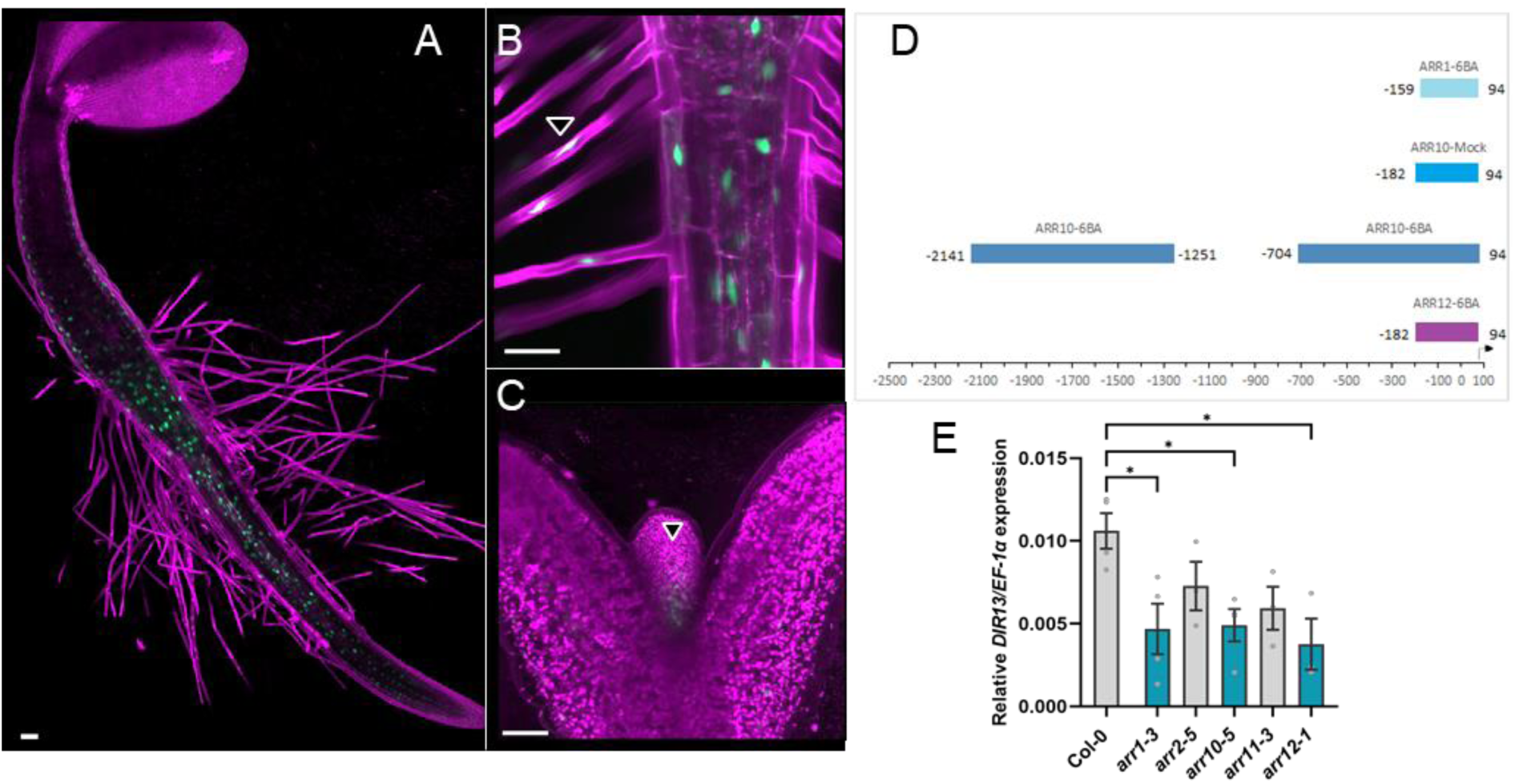
D*I*R13 promoter is active in all root tissues at early developmental stage and is the target of type-B ARR1, ARR10 and ARR12 transcription factors. **(A - C)** Activity of *DIR13* promoter in 3-day-old seedling expressing *pDIR13:NLS-3xGFP*. The fluorescent signal was detected mostly in the root (**A**), also in root hairs (**B**) and in the shoot apical meristem (**C**) as indicated by arrowheads. The plasma membrane signal from PI staining is shown in magenta and GFP is in green. Scale bars represent 50 μm. (**D**) *DIR13* promoter analysis identifies ChIP-Seq-derived binding events (from Xie et al., 2018) for type-B ARR transcription factors (TFs) involved in cytokinin signaling pathway, in particular ARR1, ARR10 and ARR12. The coordinates represent positions relative to the transcription start site (value = 0) whereas translation start site is indicated by an arrow. (**E**) *DIR13* expression is regulated by type-B ARR1, ARR10 and ARR12 TFs. The expression level of *DIR13* was examined by RT-qPCR in *arr1-3, arr2-5, arr10-5, arr11-3* and *arr12-1* single mutants. Transcript levels were normalized to the reference *EF-1α* gene and the relative *DIR13/EF-1α* expression ratio is shown. Data are the means of a representative experiment. Error bars indicate +/- SE (n = 4). Asterisks indicate statistically significant differences at *P* < 0.05 between Col-0 WT and mutant genotypes based on one-way ANOVA followed by Dunnett’s HSD test.

**Supplemental Figure 2.**
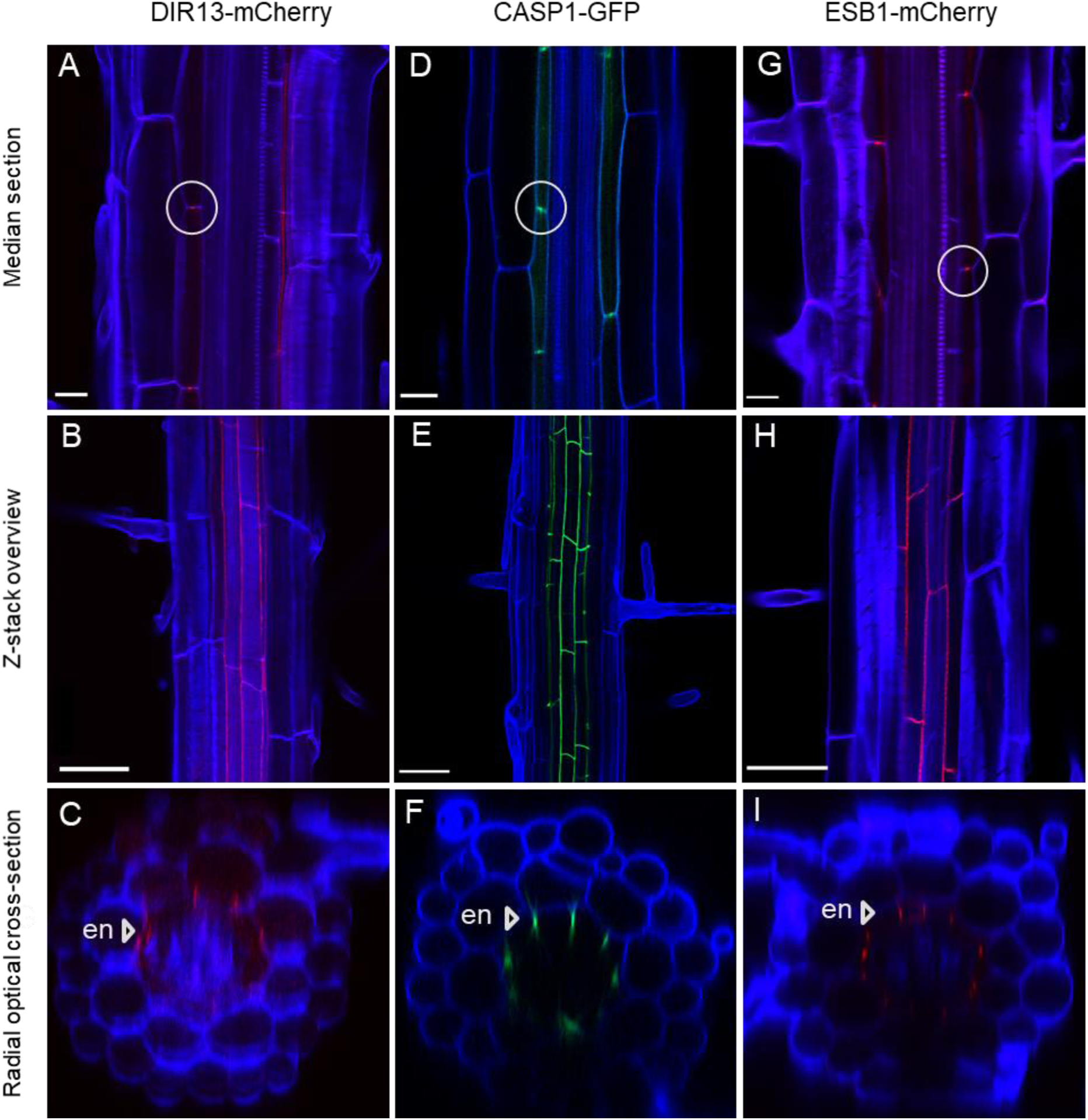
Comparison of the localization patterns of DIR13-mCherry, CASP1- GFP and ESB1-mCherry. Representative images of root differentiation zone from 7-day-old seedlings expressing *DIR13-mCherry* **(A – C),** *CASP1-GFP* **(D – F)** and *ESB1-mCherry* **(G – I)** under their native promoters. The tested reporters show similar pattern of protein localization in median sections as indicated by white circles (A, D, G), in Z-stack overview (B, E, H) and in radial optical cross-sections as indicated by arrowheads (C, F, I) respectively. Cleared roots were stained with Calcofluor White, the cell wall is shown in blue, GFP is in green, and mCherry is in red. Scale bars represent 50 μm. en, endodermis.

**Supplemental Figure 3.**
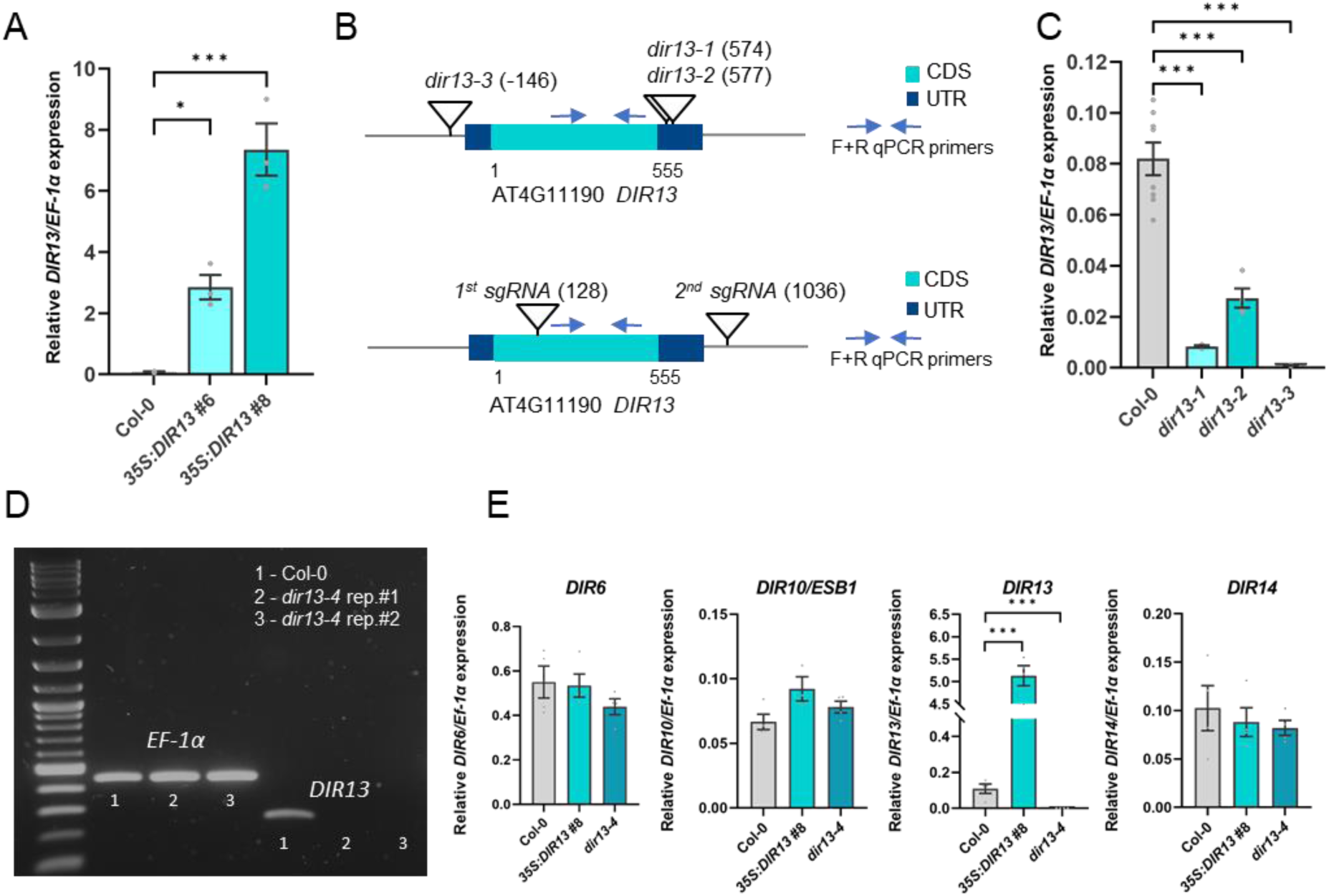
Characterization of *DIR13* overexpressing and mutant lines. **(A)** *DIR13* expression level in *DIR13* overexpressing lines. *DIR13* expression was analyzed by RT-qPCR in the roots of 11-day-old *35S:DIR13* #6 and #8 seedlings. **(B)** Schemes representing the structure of *DIR13* gene, the position of T-DNA insertions in *dir13-1*, *dir13-2* and *dir13-3* mutants (top panel), and sgRNAs used for CRISPR-Cas9 (low panel). Coding sequence (CDS) and untranslated regions (UTR) are depicted in cyan and dark blue respectively. Blue arrows indicate the positions of the primers used for RT-qPCR and semi-quantitative RT-PCR amplification of *DIR13*. **(C)** *DIR13* expression was detected by RT-qPCR in the roots of 11-day-old *dir13-1*, *dir13- 2* and *dir13-3* T-DNA insertion mutants. **(D)** Semi-quantitative RT-PCR gel electrophoresis shows the absence of *DIR13* transcripts in *dir13-4* mutant line. *EF-1α* was used as a control. **(E)** The expression levels of *DIR13* closest homologs are not affected by *DIR13* overexpression or *dir13-4* knock-out mutation. Expression levels of *DIR6, DIR10 (ESB1), DIR13 and DIR14* were verified by RT-qPCR in the roots of 11-day-old Col-0 WT, *35S:DIR13* #8 and *dir13-4* seedlings. In (A, C and E), transcript levels were normalized to the reference *EF-1α* gene and the relative *DIR* target gene to *EF-1α* expression ratios are shown. Data are the means of a representative experiment. Error bars indicate +/- SE (n ≥ 3). Asterisks indicate statistically significant differences between genotypes based on one-way ANOVA followed by Tukey’s HSD test (\**P* < 0.05, \*\*\**P* < 0.001).

**Supplemental Figure 4.**
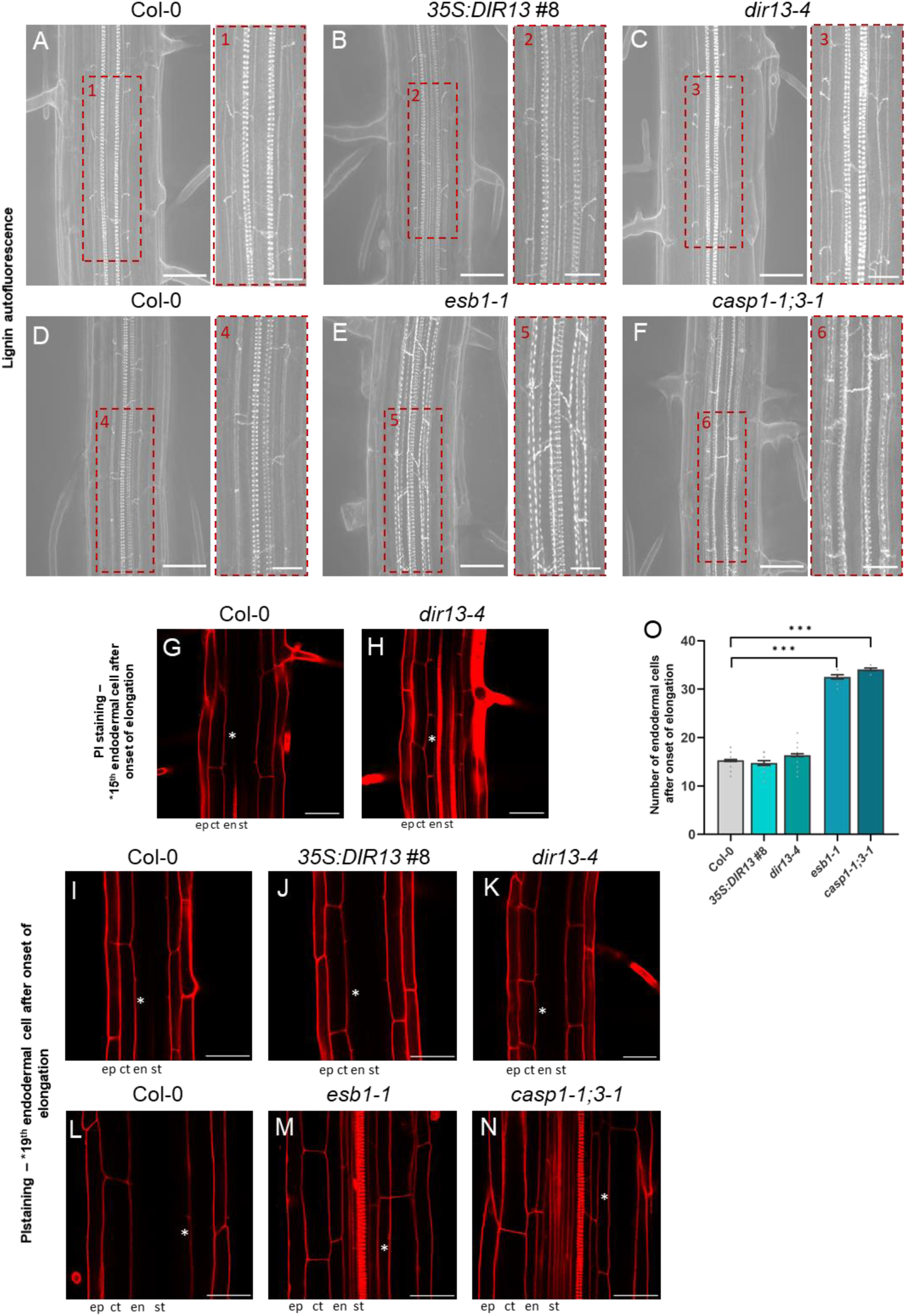
DIR13 is not required for normal morphology and function of Casparian strip. **(A – F)** Visualization of lignin autofluorescence after clearing roots of 7-day-old Col-0 WT (A, D), *35S:DIR13* #8 (B), *dir13-4* (C), *esb1-1* (E), and c*asp1-1;3-1* (F), and their close-up images (1 - 6) shown respectively in red dashed rectangles. **(G – N)** Propidium iodide (PI) penetration into the stele in 5-day-old Col-0 WT (G, I, L), *dir13-4* (H, K), *35S:DIR13* #8 (J), *esb1-1* (M) and c*asp1-1;3-1* (N). Asterisks mark the 15^th^ (G, H) and 19^th^ (I - N) endodermal cell after onset of elongation. The onset of elongation (e.g., fully expanded cell) is defined as the zone where the length of an endodermal cell was observed to be more than twice its width. Scale bars represent 50 μm in (A - O) and 20 μm in (1 - 6). ep, epidermis; ct, cortex; en, endodermis; st, stele. **(O)** Quantification of PI penetration in Col-0 WT, *35S:DIR13* #8, *dir13-4*, *esb1-1*, and *casp1-1;3-1* roots. The number of endodermal cells showing PI staining into the stele were counted from the first fully expanded endodermal cell. Data are the means of a representative experiment. Error bars indicate +/- SE, (n ≥ 10). Asterisks indicate statistically significant differences between Col-0 WT and mutant lines based on Poisson mixed model analysis and Tukey test (\*\**P* < 0.01, \*\*\**P* < 0.001).

**Supplemental Figure 5.**
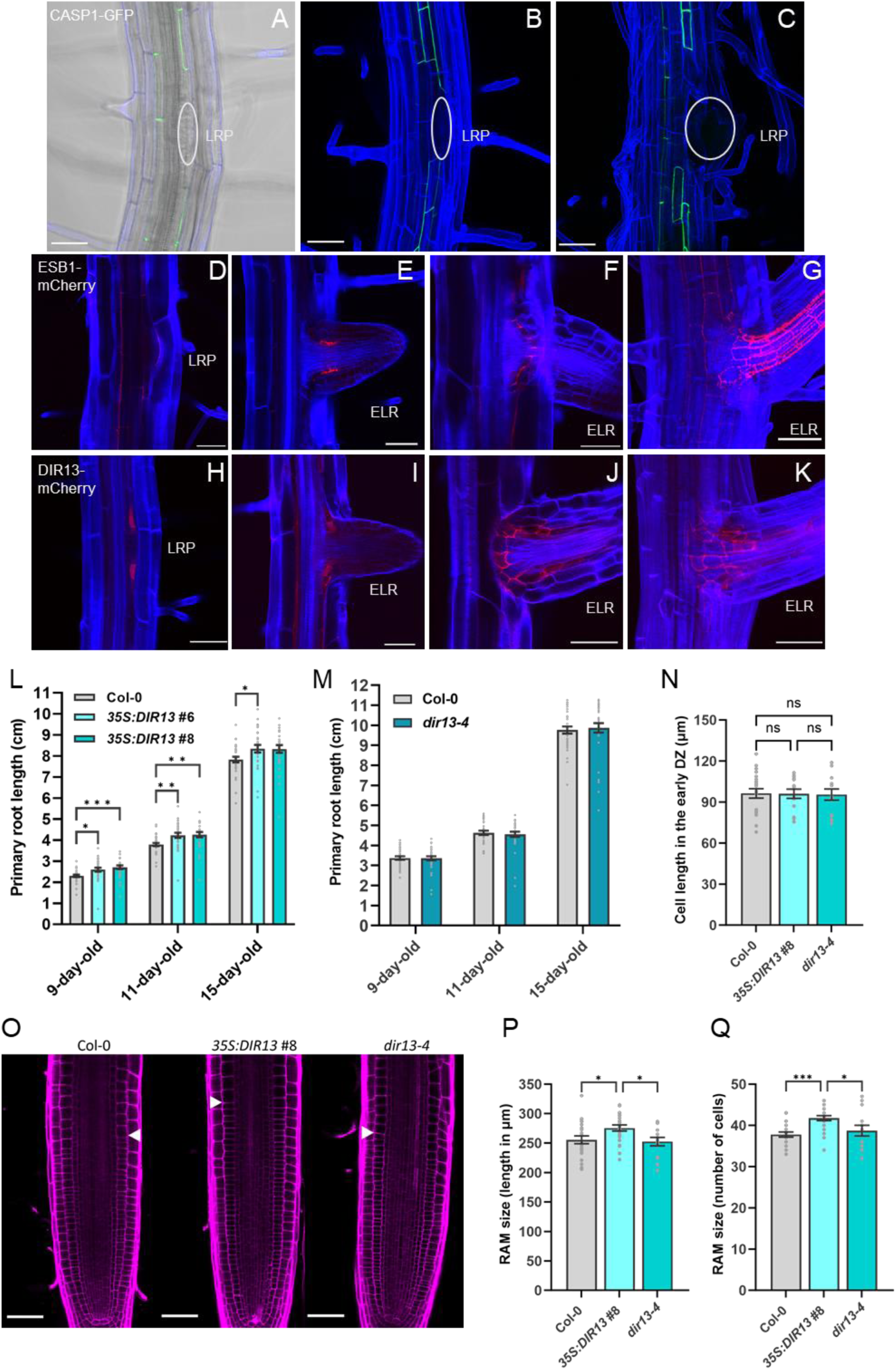
DIR13, CASP1 and ESB1 show distinct localization patterns during lateral root formation. **(A – C)** CASP1-GFP signal is not detected in lateral root primordia (LRP) as indicated by white ovals. Representative images of 9-day-old seedlings where (A) is confocal image merged with transmission image and (B, C) are Z-stack overviews. **(D – G)** ESB1-mCherry localizes in both LRP and emerged lateral root (ELR). Representative images of Z-stack overviews of 9-day-old seedlings expressing *pESB1:ESB1-mCherry* in the developing lateral root area of LRP (D) and ELR (E – G). **(H – K)** DIR13-mCherry localization partially resembles ESB1-mCherry expression pattern but is more localized to the LR peripheral basal cells throughout LR development. Representative images of Z-stack overviews of 9-day-old seedlings expressing *pDIR13:DIR13-mCherry* in the developing lateral root area of LRP (H) and ELR (I – K). Cleared roots were stained with Calcofluor White, the cell wall is shown in blue, GFP is in green and mCherry is in red. Scale bars represent 50 μm. **(L – M)** Comparison of the primary root length (in cm) in Col-0 WT, *35S:DIR13* #6 and #8 (F) and *dir13-4* mutant (G) seedlings grown vertically for 9, 11 and 15 days. Data are the means of a representative experiment. Error bars indicate +/- SE (n ≥ 30). Asterisks indicate statistically significant differences between genotypes based on two-way ANOVA followed by Dunnett’s HSD test (\**P* < 0.05, \*\**P* < 0.01, \*\*\**P* < 0.001). **(N)** Cell length measured in the early differentiation zone (DZ) of Col-0 WT, *35S:DIR13* #8 and *dir13-4* mutant. Data are presented as the mean +/-SE (n≥14). No statistically significant difference was found between tested genotypes based on one-way followed by Tukey’s HSD test. **(O)** Representative images of 7-day-old roots of Col-0 WT, *35S:DIR13* #8 and *dir13-4* mutant. The plasma membrane signal from PI staining is shown in magenta. The end of the measured RAM size indicated by white arrowheads. Scale bars represent 50 µm. **(P, Q)** RAM size measured as total length from QC until the distal end of TZ (in µm) (P) and as number of cortex cells in the same area (Q). Asterisks indicate statistically significant differences between tested genotypes based on one-way ANOVA (P) or Poisson mixed model analysis (Q) followed by Tukey’s HSD test (\**P* < 0.05, \*\*\**P* < 0.001).

**Supplemental Figure 6.**
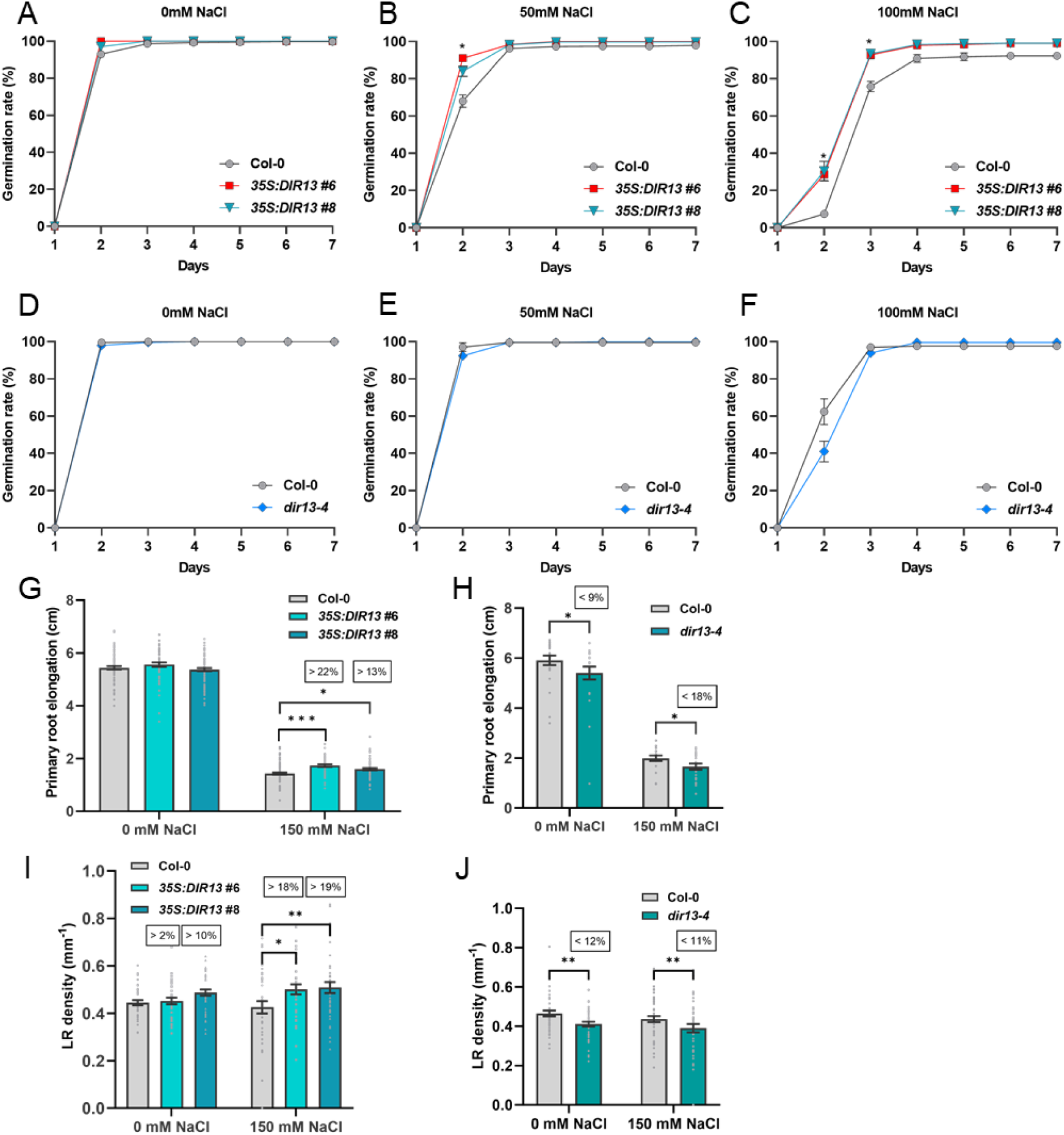
DIR13 improves tolerance to salt stress. **(A - F)** Seed germination rates of *35S:DIR13* #6 and #8 overexpressing lines (A - C) and *dir13-4* mutant (D - F) under salt stress treatment. The germination rate was recorded for 7 days after stratification in the presence of 0, 50 and 100 mM NaCl. Data are the averages of three independent experiments. Error bars indicate +/- SE (n ≥ 100). Asterisks indicate statistically significant differences at *P* < 0.05 between Col-0 WT and *DIR13* OE in (A - C) based on repeated measures two-way ANOVA followed by Dunnett’s HSD test. **(G - H)** DIR13 attenuates salt stress-mediated inhibition of primary root growth. 5-day-old Col-0 WT, *35S:DIR13* #6 and #8 lines (G) and *dir13-4* mutant (H) seedlings grown on MS/2 media were transferred to MS/2 media supplemented with 0 mM (control) or 150 mM NaCl. Primary root elongation (in cm) was measured 7 days after transfer. Data are the averages of three independent experiments. Error bars indicate +/- SE (n ≥ 60). Asterisks indicate statistically significant differences between genotypes based on two-way ANOVA followed by Tukey’s HSD test (\**P* < 0.05, \*\*\**P* < 0.001). Numbers in rectangles show the relative difference compared to Col-0 WT in percentage. **(I - J)** DIR13 promotes lateral root growth even in the presence of salt. LR density (lateral root number per mm of primary root length) was measured 9 days after the transfer of Col-0 WT, *35S:DIR13* #6 and #8 (I) and *dir13-4* (J) seedlings to 150 mM NaCl. Data show the means of a representative experiment +/- SE (n ≥ 20). Asterisks indicate statistically significant differences between genotypes based on two-way ANOVA followed by Tukey’s HSD test (\*\**P* < 0.01, \*\*\**P* < 0.001). Numbers in rectangles show the relative difference compared to Col-0 in percentage.

**Supplemental Figure 7.**
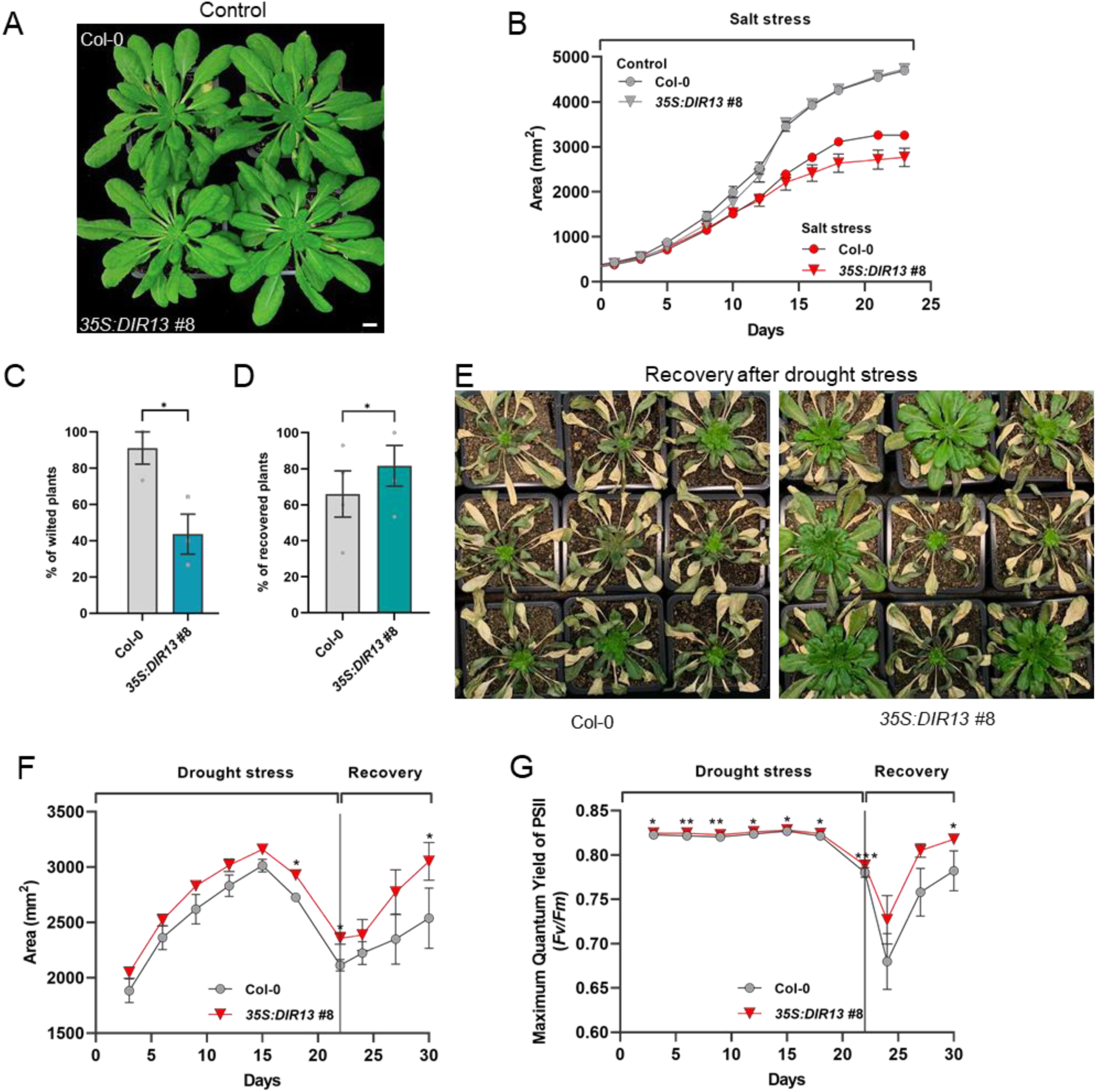
Tolerance to drought stress of *DIR13* overexpressing plants is higher than Col-0 WT. **(A)** Phenotypes of 8-week-old Col-0 WT and *35S:DIR13* #8 plants growing under normal short-day conditions. Scale bar represents 1 cm. **(B)** Area (in mm^2^) of the plant rosette (A; Fig. 4 G) was recorded in Col-0 WT and *35S:DIR13* #8 plants during the control and salt stress conditions. Data are the means of a representative experiment. Error bars indicate +/- SE (n ≥ 7). No statistically significant differences between Col-0 WT and *35S:DIR13* #8 in control or salt stress conditions was found based on mixed model analysis followed by Tukey’s HSD test. **(C, D)** *DIR13* overexpression promotes plant tolerance to drought stress. Percentage of wilted Col-0 WT and *35S:DIR13* #8 plants (B) was calculated 3 weeks after water withholding and percentage of recovered plants (C) was calculated 5 days after rewatering. The data are means of three independent experiments. Error bars indicate +/- SE (n = 45). Asterisks indicate statistically significant differences at *P* < 0.05 between Col-0 WT and *35S:DIR13* #8 based on repeated measures one-way ANOVA followed by Tukey’s HSD test. **(E)** Representative images of the recovery phenotype after 5 days of rewatering drought stressed Col-0 WT and *35S:DIR13* #8 plants. **(F, G)** Area (in mm^2^) of the plant rosette (E) and Photosystem II maximum quantum yield (*Fv*/*Fm*) efficiency (F) were recorded in Col-0 WT and *35S:DIR13* #8 plants during the drought stress and recovery phases. Data are the means of a representative experiment. Error bars indicate +/- SE (n = 13). Asterisks indicate statistically significant differences between Col- 0 WT and *35S:DIR13* #8 based on mixed model analysis followed by Tukey’s HSD test (\**P* < 0.05, \*\**P* < 0.01).

**Supplemental Figure 8.**
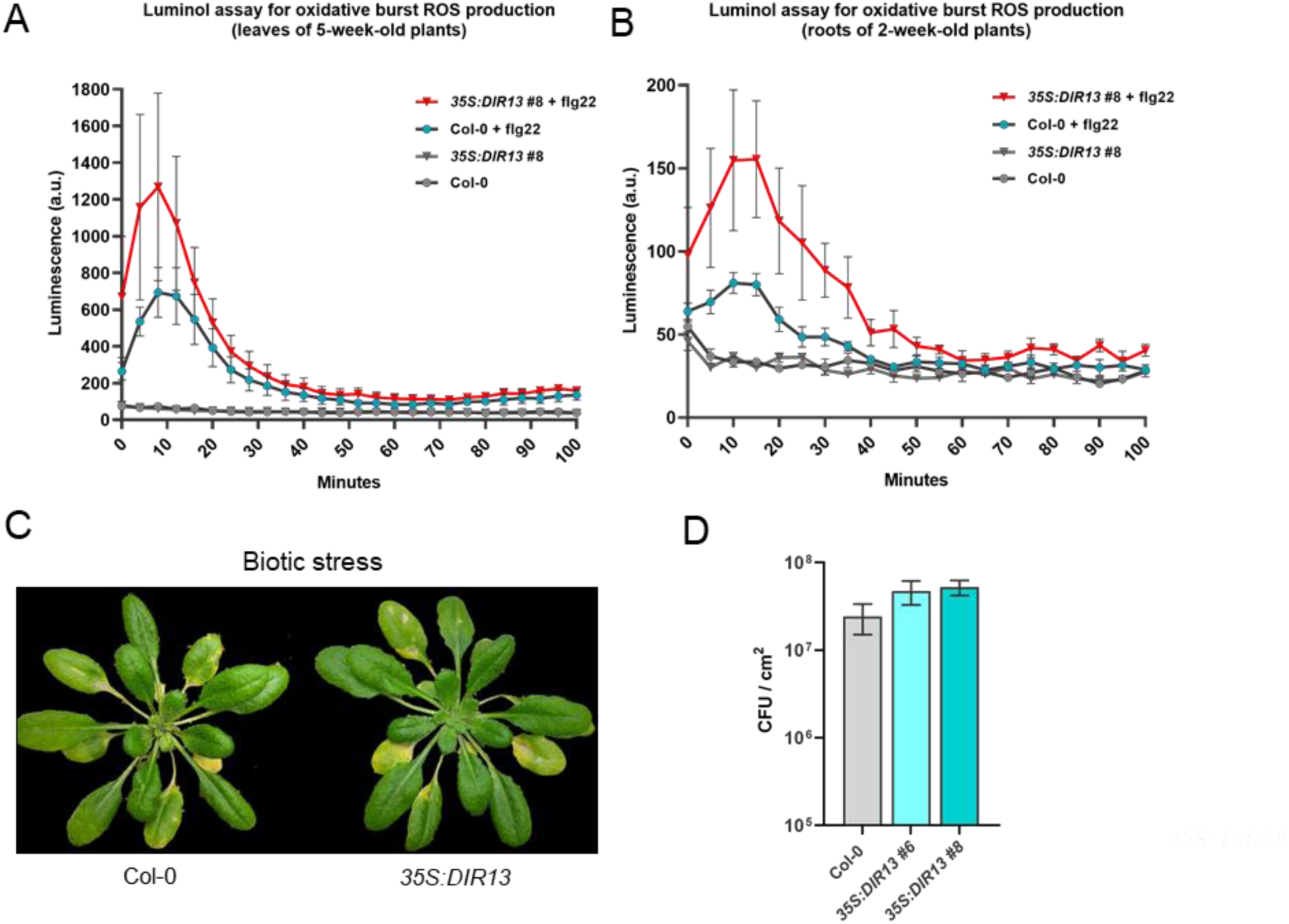
Biotic stress responses in Col-0 WT and *35S:DIR13* #8 plants. **(A, B)** *35S:DIR13* #8 plants show enhanced ROS production triggered by flg22. Oxidative burst induced by 1 µM flg22 was measured by luminol assays in 5-week-old leaves (A) and 2-week-old roots (B) of Col-0 WT and *35S:DIR13* #8 plants. Data show the means of a representative experiment. Error bars indicate +/- SE (n = 6). Statistically significant difference at *P* < 0.05 between Col-0 WT and *35S:DIR13* #8 was not found based on repeated measures mixed model analysis followed by Tukey’s HSD test. **(C)** Disease symptoms in 5-week-old Col-0 WT and *35S:DIR13* #8 plants 5 days after spray-inoculation with *Pseudomonas syringae* pv. tomato (*Pst*) DC3000 at 10^8^ cfu/mL. **(D)** Quantification of *Pst* DC3000 bacterial growth (colony forming units – CFU per cm^2^ of leaf area) in Col-0 WT and *35S:DIR13* #6, #8 leaves 3 days after spraying with 10^8^ cfu/mL *Pst* DC3000. Results are the means of two independent experiments. Error bars indicate +/- SE (n = 16). No significant difference at *P* < 0.05 between Col-0 WT and *35S:DIR13* #6, #8 was found based on repeated measures one-way ANOVA followed by Tukey’s HSD test.

**Supplemental Figure 9.**
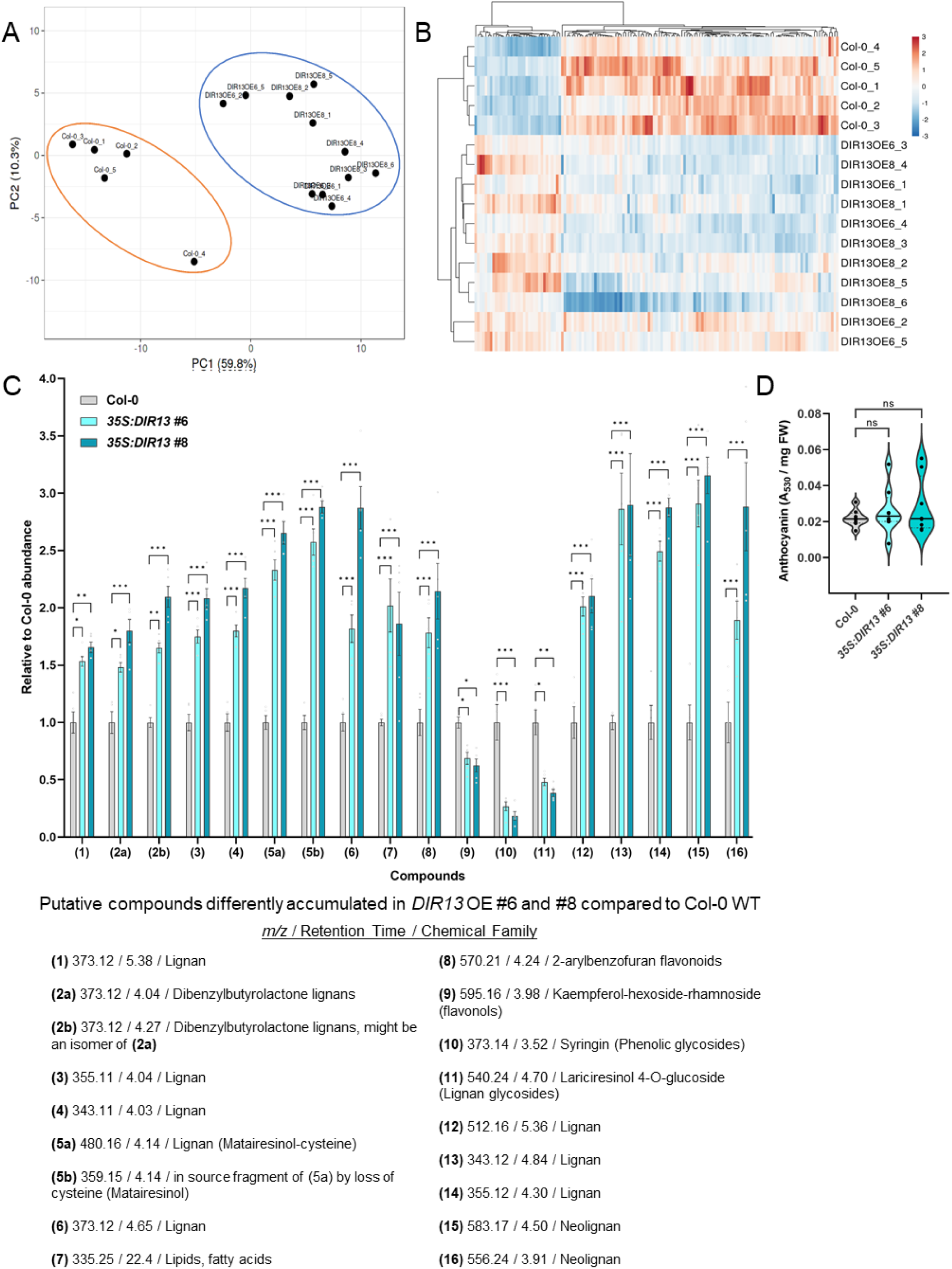
Untargeted metabolomic analyses reveal changes in the accumulation of putative lignans and neolignans in *DIR13* overexpressing lines. **(A) Principal** Component Analysis (PCA) of HPLC-MS/MS data in ElectroSpray Ionization in positive (ESI+) mode from root biological replicates of 10-day-old Col-0 WT, *35S:DIR13* #6 and #8. PCA demonstrates the difference between Col-0 WT and *DIR13* OE lines in the accumulation of significantly different abundant compounds. **(B)** Clustering of compounds detected by HPLC-MS/MS (ESI+ mode) which are significantly different abundant between Col-0 WT and *DIR13* OE roots. Columns were centered, unit variance scaling was applied to columns. Both rows and columns were clustered using correlation distance and average linkage. **(C)** *DIR13* overexpression leads to the accumulation of putative lignans and neolignans in roots. Selected differentially abundant compounds annotated mostly as putative (neo)-lignans were extracted from Col-0 WT, *35S:DIR13* #6, #8 roots and detected by HPLC-MS/MS in ESI+ mode. Data are shown as relative to Col-0 WT and are the means +/- SE (n ≥ 5). Asterisks indicate statistically significant differences between Col-0 WT and *35S:DIR13* #6, #8 based on two-way ANOVA followed by Dunnett’s HSD test. (\**P* < 0.05, \*\**P* < 0.01, \*\*\**P* < 0.001). **(D)** Quantification of anthocyanins Abs_530_/mg of fresh weight (FW) in the shoots of 11-day-old Col-0 WT and *35S:DIR13* #6 and #8 seedlings. No significant difference at *P* < 0.05 between Col-0 WT and *35S:DIR13* #6, #8 was found based on one-way ANOVA followed by Tukey’s HSD test.

**Supplemental Figure 10.**
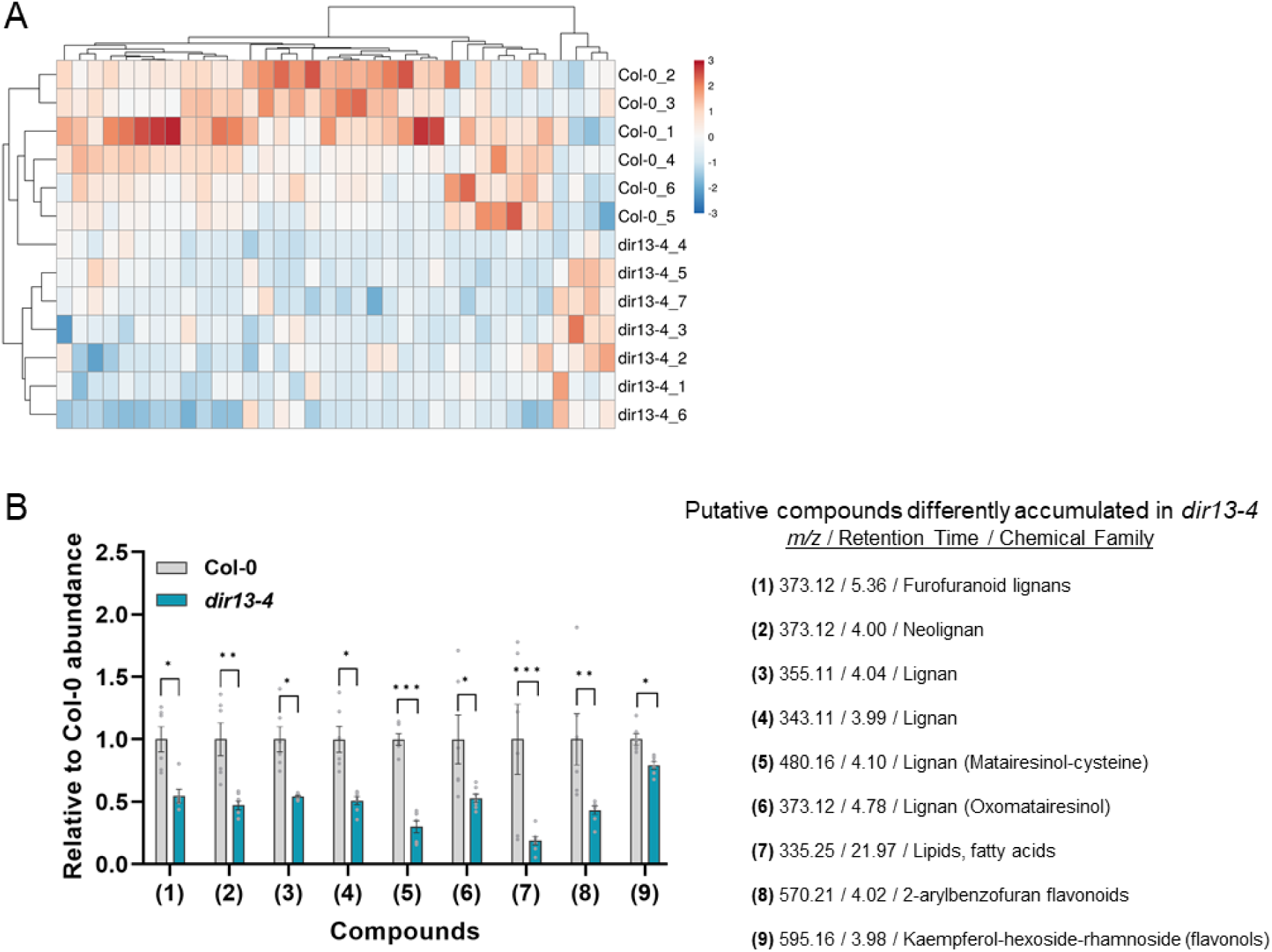
Untargeted metabolomic analyses reveal changes in the accumulation of putative lignans and neolignans in *dir13-4* mutant line in ESI+ mode. **(A)** Clustering of compounds detected by HPLC-MS/MS which are significantly different presented in Col-0 WT and *dir13-4* plants. Columns are centered, unit variance scaling is applied to columns. Both rows and columns are clustered using correlation distance and average linkage. **(B)** Differential accumulation of phenolic compounds in *dir13-4* mutant. Selected differentially abundant compounds annotated mostly as putative (neo)-lignans were extracted from Col-0 WT, *dir13-4* roots and detected by HPLC-MS/MS in ESI+ mode. Data are shown as relative to Col-0 WT and are the means +/- SE (n = 6). Asterisks indicate statistically significant differences between Col-0 WT and *dir13-4* based on two-way ANOVA followed by Tukey’s HSD test. (\**P* < 0.05, \*\**P* < 0.01, \*\*\**P* < 0.001).

**Supplemental Figure 11.**
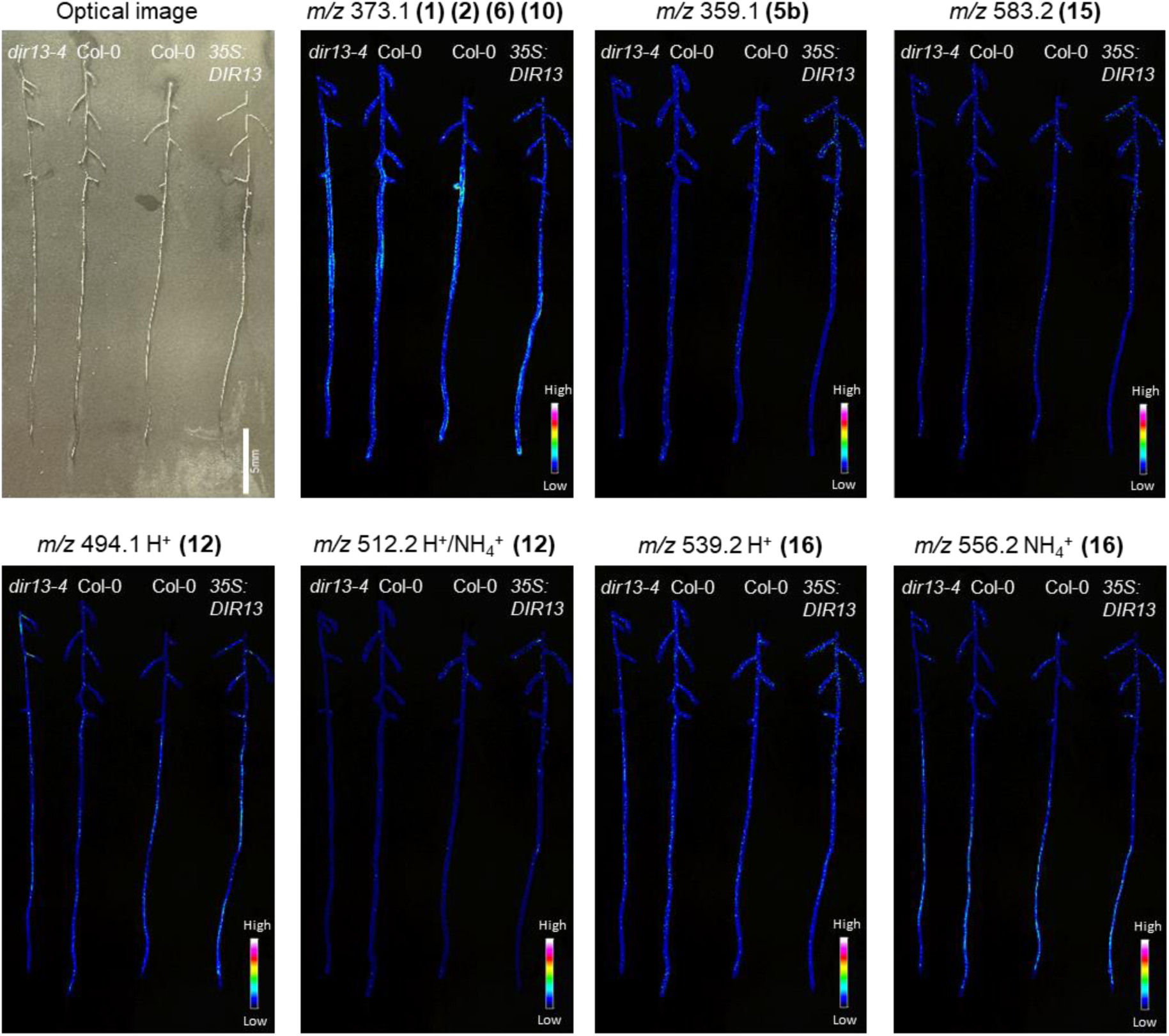
Spatial distribution of putative lignans in the roots of *dir13-4* mutant and *DIR13* overexpressing lines. Imaging Mass Spectrometry (IMS) of roots from 10- day-old Col-0 WT, *dir13-4* mutant and *35S:DIR13* #8 overexpressing lines. Signals at *m/z* 373.1 (1) (2) (6) (10), *m/z* 359.1 (5b), 583.2 (15), 494.1 H^+^ (12), 512.2 H^+^/NH_4_^+^ (12), 539.2 H^+^ (16) and 556.2 NH_4_^+^ (16) correspond to putative matairesinol (5b), oxomatairesinol (6), syringin (10) and metabolites annotated as lignans (1), (12) or neolignans (2), (15) and (16). For metabolites (12) and (16) the *m/z* signals from two possible ionizations by H^+^ or NH_4_^+^ are shown. The spatial resolution is 50 µm. MS signal intensity is indicated by color bars and the scale bar represents 5 mm. The experiment was repeated at least twice with similar results.

**Supplemental Figure 12.**
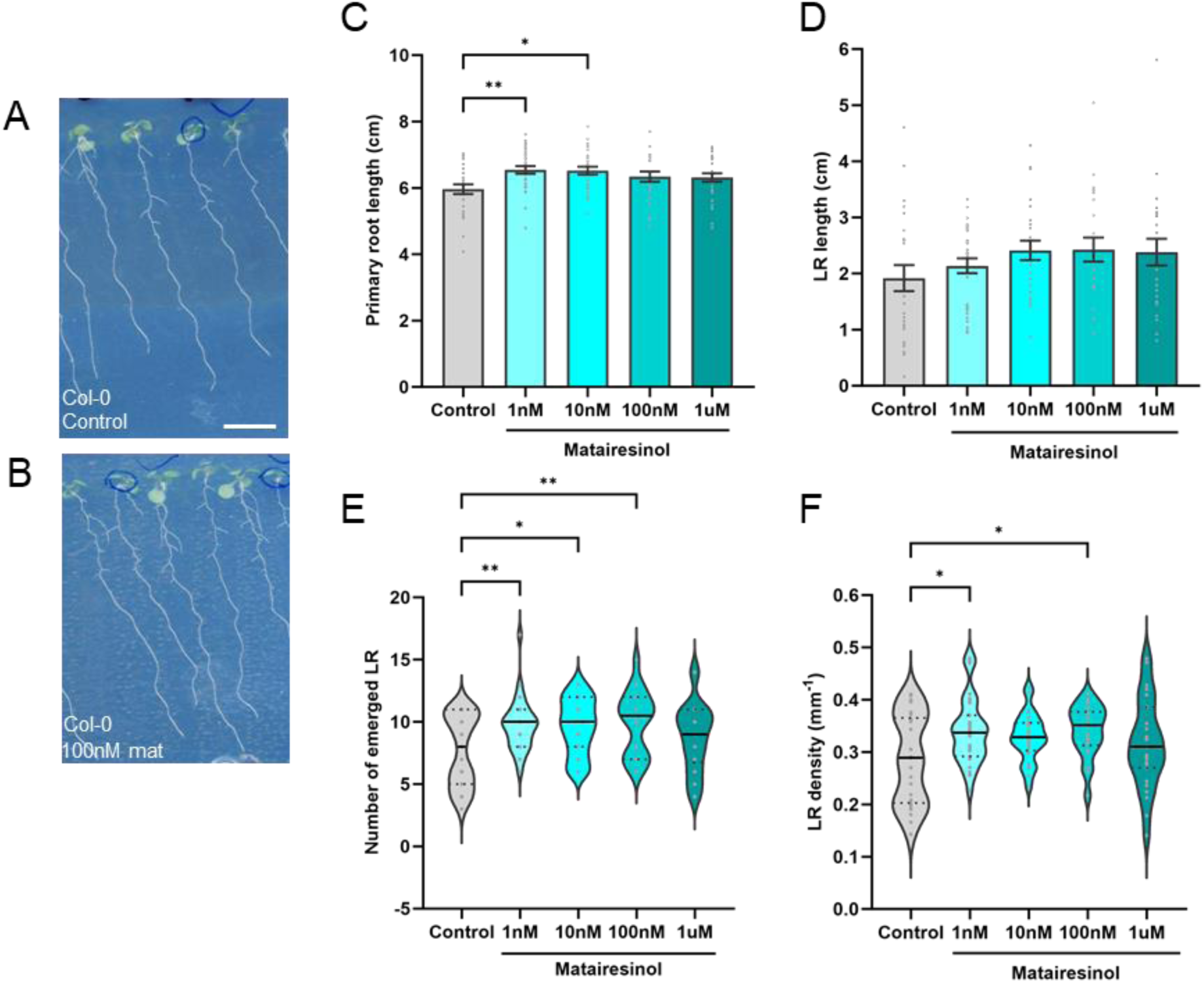
Matairesinol promotes lateral root growth. **(A - B)** Phenotype of 11- day-old Col-0 WT seedlings growing in normal conditions (A) or on plates supplemented with 100nM matairesinol (B). Scale bars represent 1 cm. **(C - D)** Comparison of the primary root length (in cm) (C) and total lateral root length per seedling (in cm) (D) in 11-day-old Col-0 WT seedlings. Data are the means of a representative experiment. Error bars indicate +/- SE (n ≥ 22). Asterisks indicate statistically significant differences between genotypes at \*\**P* < 0.01 based on one-way ANOVA followed by Dunnett’s HSD test. **(E - F)** Average number of emerged lateral roots (E) and LR density expressed as the number of lateral roots per cm of primary root length (F) in 11-day-old Col-0 WT seedlings. Data show the median and upper and lower quartiles of a representative experiment (n ≥ 22). Asterisks indicate statistically significant differences between genotypes at \**P* < 0.05 based on Poisson mixed model analysis (E), one-way ANOVA followed by Dunnett’s HSD test (F).

**Supplemental Figure 13.**
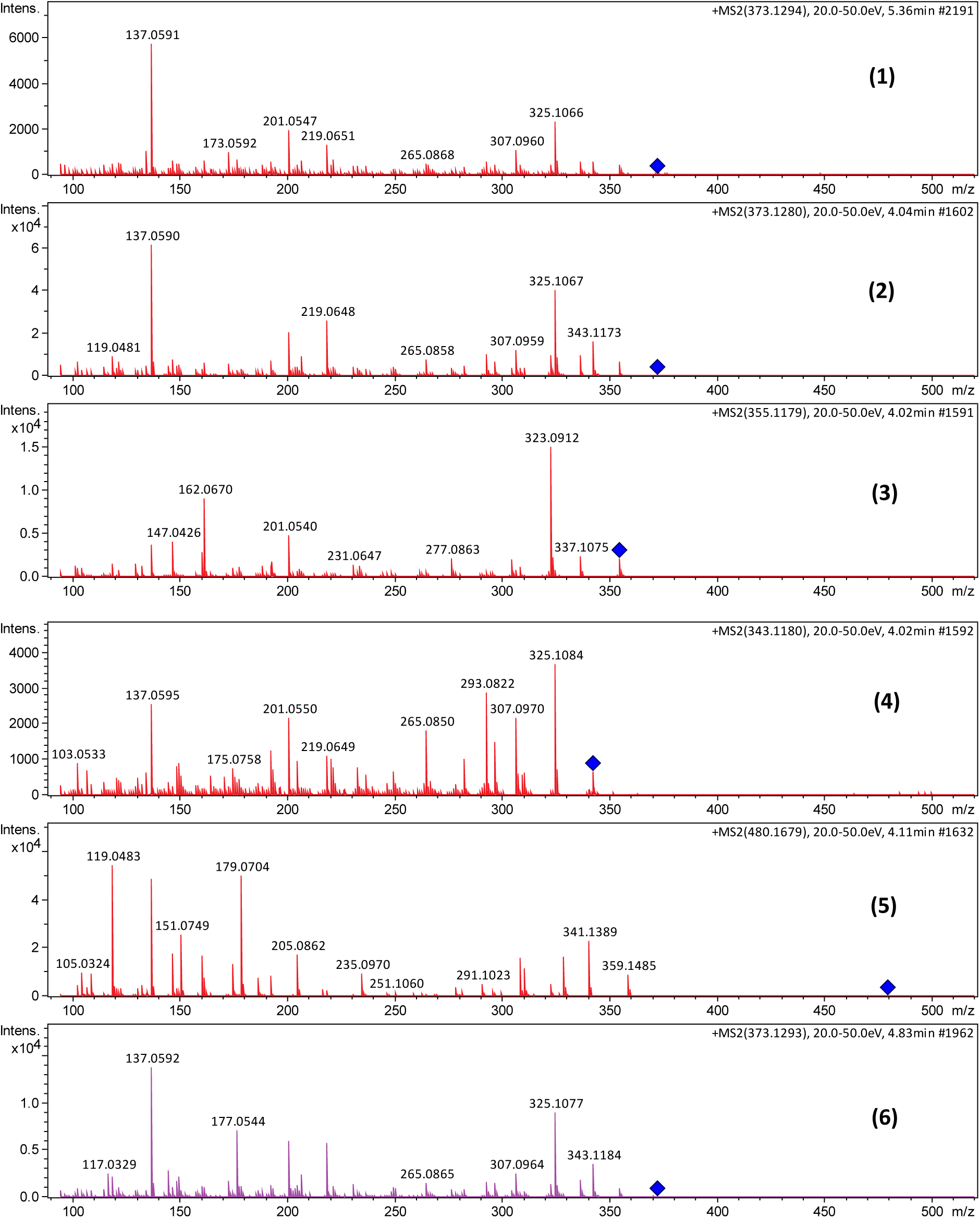
ESI+ MS/MS spectra of accumulated putative lignans and neolignans in roots overexpressing *DIR13*. For the compounds numbering see Figure 6B, HPLC-MS/MS analyses were performed using *35S:DIR13* #8.

## Supplemental Materials and Methods

### Lignin autofluorescence

Lignin autofluorescence experiment was performed as described by (Hosmani et al., 2013). 5- to 6-day-old seedlings were transferred to a Petri dish containing 0.24 M HCl and 20 % methanol and incubated for 15 minutes at 57 °C. The solution was replaced with 7 % NaOH and 60 % ethanol and incubated for 15 minutes at room temperature. After these clearing steps, roots were rehydrated at room temperature by sequential replacement of the solution using 40 % ethanol for 5 minutes, 20 % ethanol for 5 minutes, 10 % ethanol for 5 minutes, 5 % ethanol for 15 minutes, and finally 25 % glycerol was added. For mounting root on glass slides, 50 % glycerol was used. Autofluorescence of the Casparian strip was observed on cleared roots (Malamy & Benfey, 1997) by using a Zeiss LSM880 confocal microscope with excitation at 488 nm and with band path filter of 500–600 nm.

### PI staining and quantification

For assay of the apoplastic barrier, PI staining was done as described previously (Alassimone et al., 2010). Briefly, 5- to 6-day-old Col-0 WT, *35S:DIR13, dir13-4, esb1-1, and casp1-1;3-1* seedlings were incubated in the dark for 10 min in a fresh solution of 15 μM (10 μg/mL) PI and rinsed twice in water. For quantification, the “onset of elongation” was defined as the point where an endodermal cell in a median optical section was more than twice its width and the number of cells were counted until PI could not penetrate into the stele.

### ClearSee-adapted cell wall staining

ClearSee-adapted cell wall staining was performed as described before by (Ursache et al., 2018). Briefly, 5- to 6-day-old seedlings were fixed in 3 ml 1 × PBS containing 4 % paraformaldehyde for 1 hour at room temperature in 6-well plates and washed twice with 3 ml 1 × PBS. Following fixation, the seedlings were cleared in 3 ml ClearSee solution under gentle shaking. After overnight clearing, the solution was exchanged to new ClearSee solution containing 0.1% Calcofluor White for cell wall staining. The dye solution was removed after 1 hour of staining and rinsed once with fresh ClearSee solution. The samples were washed in new ClearSee solution for 30 minutes with gentle shaking and washed again in another fresh ClearSee solution for at least 1 hour before observation.

### Drought stress tolerance tests

For the drought tolerance test, different genotypes randomly mixed and initially grown on soil under normal watering conditions for 4 weeks were subjected to a 21-day dehydration period by withholding water, during which observations were made. The percentage of wilted plants was counted when wild-type plants exhibited wilting on young/meristematic leaves. Subsequently, watering was resumed, and the percentage of recovered plants was calculated 5 days after rewatering and counted as plants with newly green meristematic leaves. Control plants grown in parallel under well-watered conditions were used as a control. During the drought experiment, the morphologic and fluorescent parameters were monitored using the automatic phenotyping platform (PlantScreen^TM^ SC Root System, PSI). Particularly, measurements of “Area (mm^2^)” and “Maximum quantum yield of PSII (*Fv/Fm*)” representing the maximal photochemical efficiency were taken every 3 days for each plant.

### Luminol assay

Apoplastic oxidative burst evaluation ROS released by leaf and root tissue was assayed as described by (Arnaud et al., 2023). 6 mm leaf discs of 4.5-week-old Arabidopsis plants were cut in 4 equal pieces or 2-week-old roots were collected from 16 seedlings for each genotype and transferred into the wells of 96-well plate containing distilled water. The plates were incubated for 3 hours at room temperature for recovery after wounding. Then water was exchanged with a solution containing 20 µM luminol (Sigma) and 10 µg/mL horseradish peroxidase (Sigma). After adding water as a control solution or 1 µM flg22, the luminescence was measured immediately using a PROMEGA microplate reader (BMG Labtech) with a reading time of 2 sec.

### Bacterial infection assay

Bacterial infection assay was performed as previously described by (Arnaud et al., 2017). Bacterial strain *Pseudomonas syringae* pv. tomato DC3000 (Pst DC3000) was cultivated overnight at 28°C in King’s B medium supplemented with rifampicin (100 µg/ml) and kanamycin (50 µg/ml) antibiotics. Bacteria were collected by centrifugation at 3000 *g* for 5 min at room temperature and washed twice in 10 mM MgCl_2_. Plants were surface inoculated by spraying with a bacterial solution of 1x10^8^ colony-forming units (cfu)/mL in 10 mM MgCl2 containing 0.02 % Silwet L-77, and plants were covered to maintain high humidity until disease symptoms developed. After 3 days, bacterial growth in the apoplast of three leaves per plant and four plants per genotype was determined as previously described (Katagiri et al., 2002).

